# A novel Na_v_1.5-dependent feedback mechanism driving glycolytic acidification in breast cancer metastasis

**DOI:** 10.1101/2023.06.16.545273

**Authors:** Theresa K Leslie, Aurelien Tripp, Andrew D James, Scott P Fraser, Michaela Nelson, Nattanan Sajjaboontawee, Michael Toss, Wakkas Fadhil, Samantha C Salvage, Mar Arias Garcia, Melina Beykou, Emad Rakha, Valerie Speirs, Chris Bakal, George Poulogiannis, Mustafa B A Djamgoz, Antony P Jackson, Hugh R Matthews, Christopher L-H Huang, Andrew N Holding, Sangeeta Chawla, William J Brackenbury

## Abstract

Solid tumours have abnormally high intracellular [Na^+^]. The activity of various Na^+^ channels may underlie this Na^+^ accumulation. Voltage-gated Na^+^ channels (VGSCs) have been shown to be functionally active in cancer cell lines, where they promote invasion. However, the mechanisms involved, and clinical relevance, are incompletely understood. Here, we show that protein expression of the Na_v_1.5 VGSC subtype strongly correlates with increased metastasis and shortened cancer-specific survival in breast cancer patients. In addition, VGSCs are functionally active in patient-derived breast tumour cells, cell lines, and cancer-associated fibroblasts. Knock down of Na_v_1.5 in a mouse model of breast cancer suppresses expression of invasion-regulating genes. Na_v_1.5 activity increases glycolysis in breast cancer cells, likely by up-regulating activity of the Na^+^/K^+^ ATPase, thus promoting H^+^ production and extracellular acidification. The pH of murine xenograft tumours is lower at the periphery than in the core, in regions of higher proliferation and lower apoptosis. In turn, acidic extracellular pH elevates persistent Na^+^ influx through Na_v_1.5 into breast cancer cells. Together, these findings show positive feedback between extracellular acidification and movement of Na^+^ into cancer cells which can facilitate invasion. These results highlight the clinical significance of Na_v_1.5 activity as a potentiator of breast cancer metastasis and provide further evidence supporting the use of VGSC inhibitors in cancer treatment.

## Introduction

Breast cancer is the leading cause of cancer-related deaths in women worldwide (1) and most deaths are due to metastatic disease resulting from poor treatment options and therapy resistance (2). Around 20-30% of patients with primary breast cancer will go on to develop distant metastasis and once this has been diagnosed, there is currently no cure available. Thus, there is an urgent need for improved treatments to prevent or reduce breast cancer metastasis.

Increasing evidence points to ion channels as key regulators of cancer progression (3–6). Members of the voltage-gated Na^+^ channel (VGSC) family are upregulated in multiple cancer types (7). In solid cancers, including breast, prostate, lung, and colon cancer, VGSC activity promotes cellular invasion (8, 9). In breast cancer, the Na_v_1.5 subtype is upregulated at the mRNA level compared to normal tissue and is associated with recurrence and metastasis (10). Na_v_1.5 is also upregulated in breast cancers at the protein level (11, 12), predominantly in its neonatal D1:S3 splice form (13); however, the sample sizes of these studies were too small to reliably determine the relationship between Na_v_1.5 expression and clinical outcome. Electrophysiological methods have not yet been used to investigate functional Na_v_1.5 activity in breast cancer tissue or primary cell cultures. Nonetheless, Na^+^ currents carried by Na_v_1.5 have been detected in a small number of breast cancer cell lines and in tissue slices from murine tumour xenografts (11, 12, 14, 15). In these cells, the persistent Na^+^ current (as distinct from the transient, inactivating Na^+^ current), which passes through the channels at the resting membrane potential (V_m_), has been shown to potentiate cellular invasion *in vitro* and tumour growth and metastasis *in vivo* (10–12, 14, 16, 17). Importantly, the metastasis-promoting function of Na_v_1.5 can be inhibited in preclinical models using VGSC blockers, including phenytoin and ranolazine, suggesting that Na_v_1.5 may represent a novel anti-metastatic target for therapeutic intervention (17, 18). Furthermore, perioperative administration of the VGSC blocker lidocaine has recently been shown to significantly improve disease-free survival in women with early breast cancer (19).

The mechanism by which VGSCs increase invasion of cancer cells is incompletely understood (8). The inward Na^+^ gradient created by the Na^+^/K^+^ ATPase (NKA), a major consumer of cellular ATP (20, 21), is used to power many important functions such as nutrient import and pH regulation (22). Thus, it would seem wasteful for cancer cells to deplete this inward gradient via Na_v_1.5 up-regulation. However, it has been shown that Na^+^ influx via Na_v_1.5 leads to extracellular acidification via the Na^+^/H^+^ exchanger, NHE1, activating pH-dependent cathepsins and promoting invasion (17, 23, 24). Because an increase in cytosolic [Na^+^] reduces the Na^+^ electrochemical gradient powering H^+^ extrusion via NHE1, the effect of Na_v_1.5 on NHE1 cannot be explained by physical means so an allosteric interaction between the two transporters has been proposed to explain the Na_v_1.5-dependent increase in H^+^ extrusion by NHE1 (23). An alternative possibility is that Na^+^ influx through Na_v_1.5, rather than the Na_v_1.5 protein itself, is responsible indirectly for increasing H^+^ extrusion through NHE1 and other pH regulators.

In this study, we aimed to delineate the relationship between Na_v_1.5 protein expression and clinical outcome in a large cohort of breast cancer patients. We record Na^+^ currents from patient tissue samples and primary cell cultures for the first time. We also sought to understand the mechanism by which Na_v_1.5 promotes invasion through studying the relationship between channel activity and extracellular acidification.

## Materials and Methods

### Ethical approvals

All animal procedures were carried out after approval by the University of York Animal Welfare and Ethical Review Body and under authority of a UK Home Office Project Licence and associated Personal Licences. Human tissue experiments were conducted in accordance with the ethical standards of the Declaration of Helsinki and according to national and international guidelines and were approved by the University of York Ethical Review Process.

### Breast cancer cell lines

MDA-MD-231, MCF7, T47D, MDA-MB-453, CAL51, BT549 and Hs578T cells were cultured in Dulbecco’s modified eagle medium (DMEM) supplemented with 5 % foetal bovine serum (FBS) and 4 mM L-glutamine (25). Molecular identity was confirmed by short tandem repeat analysis (26). MDA-MB-231 cells stably expressing shRNA targeting *SCN5A* were maintained in medium containing G418 (400 μg/ml) (11). LS11-083 hTERT-immortalized primary breast cancer associated fibroblast cells were also cultured in DMEM with 5 % FBS and 4 mM L-glutamine (27). Cultures were confirmed to be *Mycoplasma*-free using the 4′,6-diamidino-2-phenylindole (DAPI) method (28).

### Orthotopic xenograft breast cancer model and tissue slice preparation

*Rag2*^−/−^ *Il2rg*^−/−^ mice were bred in-house and females over the age of 6 weeks were used for tumour implantation. A suspension of 1 x 10^6^ MDA-MB-231 cells in Matrigel (Corning; 50% v/v in phosphate-buffered saline (PBS)) was implanted into the left inguinal mammary fat pad of each animal whilst under isoflurane anaesthesia. Mice were weighed and their body condition and tumour size were checked at least every 2 days. Tumours were measured using callipers and the tumour volume was calculated using the modified ellipsoidal formula, volume = ½(length × width^2^). Mice were euthanized after approximately four weeks. Tumours were dissected immediately after euthanasia and sliced in ice-cold PBS using a Campden 5100MZ vibratome to a thickness of 200 μm for patch clamp recording or 500 μm for pH-sensitive microelectrode recording.

### Human breast cancer tissue and primary cells

Biopsy samples from four patient breast tumours that were excess to pathology requirements were acquired via the Breast Cancer Now Tissue Bank (BCNTB). The samples were transported in culture medium on ice and arrived within 24 h of surgical resection. Fresh tissue slices (250 μm thick) were cut in ice-cold PBS using a vibratome (Campden 5100MZ). For isolation of single cells from the tumour tissue, fragments were cut and then dissociated using a MACS Tumour Dissociation Kit (Miltenyi Biotec). Primary cultures of breast cancer cells, enriched for carcinoma cells, and primary cultures of purified normal breast epithelial cells were also acquired via the BCNTB. Human primary cells and tumour slices were cultured in DMEM:F12 with 1 ml/100 ml penicillin/streptomycin, 2.5 μg/ml Fungizone, 10 % FBS, 0.5 μg/ml hydrocortisone, 10 μg/ml apo-transferrin, 10 ng/ml human EGF and 5 μg/ml insulin. The cells were cultured on collagen-coated glass coverslips in plastic dishes.

### Whole-cell patch clamp recording

The whole-cell patch clamp technique was used to record cell membrane currents from cells grown on glass coverslips (29). Filamented borosilicate capillary tubes were pulled and fire-polished to a resistance of ∼5 MΩ for recording from cell lines and ∼10 MΩ for recording from primary cells. The extracellular physiological saline solution (PSS) contained (in mM) NaCl 144, KCl 5.4, MgCl_2_ 1, CaCl_2_ 2.5, HEPES 5, D-glucose 5.6, and was adjusted to pH 7.2 (unless otherwise stated) using NaOH. The intracellular recording solution for measuring Na^+^ currents contained (in mM) NaCl 5, CsCl 145, MgCl_2_ 2, CaCl_2_ 1, HEPES 10, EGTA 11 and was adjusted to pH 7.4 (unless otherwise stated) using CsOH (29). The intracellular recording solution for measuring K^+^ currents contained (in mM) NaCl 5, KCl 145, MgCl_2_ 2, CaCl_2_ 1, HEPES 10, EGTA 11 and was adjusted to pH 7.4 using KOH. Recordings were made using a MultiClamp 700B amplifier (Molecular Devices). Currents were digitized using a Digidata 1440A interface (Molecular Devices), low pass filtered at 10 kHz, sampled at 50 kHz, and analysed using pCLAMP 10.7 software (Molecular Devices). For detection of small currents in primary cells, patient tumour slices and a panel of cell lines, signals were post-filtered at 1 kHz. For examination of pH dependency in MDA-MB-231 cells, currents were noise-corrected by subtracting half the peak-to-peak noise measured during the 10 ms period before depolarisation (29). Series resistance was compensated by 40-60% and linear components of leak were subtracted using a P/6 protocol (30). Cells were clamped at a holding potential of - 120 mV for 250 ms. Two main voltage clamp protocols were used, as follows:

1. To assess the voltage-dependence of activation of VGSCs and K^+^ channels, cells were held at −120 mV for 250 ms and then depolarised to test potentials in 5-10 mV steps between −120 mV and +30 mV for 50 ms.
2. To assess the voltage-dependence of steady-state inactivation, cells were held at −120 mV for 250 ms followed by prepulses for 250 ms in 5-10 mV steps between −120 mV and +30 mV and a test pulse to −10 mV for 50 ms.

### Pharmacology

Tetrodotoxin citrate (TTX, HelloBio HB1035) was diluted in sterile-filtered water to a stock concentration of 1 mM TTX/8.44 mM citrate and stored at −30 °C. The working concentration was 30 μM TTX/253 μM citrate. Ouabain octahydrate (Sigma O3125) was diluted in DMSO to a stock concentration of 50 mM and stored at −30 °C. The working concentration was 300 nM. Cariporide (Santa Cruz Biotechnology SC337619) was diluted in DMSO to a stock concentration of 50 mM and stored at - 30 °C. The working concentration was 20 μM. Sodium iodoacetate (Acros Organics 170970250) was diluted in water and made up immediately before each experiment. The working concentration was 2 μM. Oligomycin (Santa Cruz Biotechnology SC201551) was diluted to a stock concentration of 10 mM in DMSO and stored at - 30 °C. The working concentration was 1 μM. The background concentration of DMSO in these solutions was ≤0.01%.

### RNA sequencing

Mice were housed (up to 4 per cage) and were chosen at random for cell implantation ensuring that both cell types were represented within each cage/block. RNA was extracted from 12 xenograft tumours (6 control MDA-MB-231 tumours, and 6 Na_v_1.5-knock-down MDA-MB-231 tumours) tissue using TRIzol (Invitrogen), according to the manufacturer’s instructions. RNA quality assessment, library preparation, and 150 bp short-read, paired-end sequencing were conducted by Novogene Europe (Cambridge UK) with 1 µg of total RNA used for sequencing library construction with NEBNext Ultra RNA Library Prep Kit for Illumina (NEB, USA). Raw FastQ files were mapped using Bbsplit from the Bbtools 39.01 suite against Mm10 and hg38 (31) and ambiguous reads were excluded. The disambiguated hg38 reads were aligned and count matrix created using Rsubread v2.12.3 (32). Differential gene expression was calculated using DESeq2 v2_1.38.1 (33). Gene set enrichment analysis (GSEA) was performed using the implementation in VULCAN v1.20.0 (34). All code for the analysis is available from https://github.com/andrewholding/RNASeq-SCN5A.

### pH-selective microelectrodes

Unfilamented borosilicate capillary tubes were pulled to a resistance of ∼5 MΩ (measured after silanization and when filled with PSS and in a recording bath). Silanization was performed at 200 °C for 15 min with N,N-dimethyltrimethylsilyamine. Microelectrodes were back filled with the following solution (in mM): NaCl 100, HEPES 20, NaOH 10, adjusted to pH 7.5. Microelectrodes were then front-filled with H^+^ ionophore I – cocktail A (Sigma) (35). Recordings were made using a MultiClamp 900A amplifier (Molecular Devices) linked to a computer running MultiClamp 900A Commander software (Molecular Devices). The headstage amplifier was a high impedance 0.0001MU Axon HS-2 (Molecular Devices). Currents were digitized using a ITC018 A/D converter (HEKA Instruments), regular oscillatory noise was reduced with a HumBug noise eliminator (Quest Scientific) and the voltage signal was low-pass filtered at 10 Hz. Voltage was recorded using Axograph software (version 1.7.6). Electrodes were calibrated and offset from junction potentials empirically measured every 12 measurements to avoid drift with repeated electrode placement. A straight line was fitted to the offset-corrected voltage/pH calibration points, and the equation of this straight line was used to calculate the corresponding tissue slice pH. Tissue slices were maintained at 30 °C and 100 % humidity in the recording chamber at the interface between air and perfused PSS. Measurements were made on the top surface of tumour tissue slices within one hour of euthanasia. In total, 12 measurements were made from each region of the slice, alternating between regions, with calibrations and bath measurements taken before, half-way through and at the end of the series of measurements.

### Immunohistochemistry

Tissue cryopreservation, sectioning and immunohistochemistry were performed as described previously (11). The following primary antibodies were used: rabbit anti active caspase 3 (1:200; R&D Systems AF835), and rabbit anti-Ki67 (1:5000; Abcam AB15580). The secondary antibody was Alexa-568 conjugated goat anti-rabbit (1:500; Invitrogen A11036). Sections were mounted in Prolong Gold + DAPI (Thermofisher). Stained sections were imaged on a Zeiss AxioScan.Z1 slide scanner at 20X. Images were viewed using Zen 3.4 (blue edition) software (Zeiss) and the maximal intensity in the red and blue (DAPI) channels were changed to maximise the visibility of positively stained cells. The minimum intensity was not changed. Images were then converted from .czi format to 8-bit .tif format. In ImageJ, a whole section was viewed at a time and the shape was matched to the drawing of the tissue slice during pH-selective microelectrode recording. Regions of interest (ROIs; 1000 x 1000 pixels) were chosen in both “core” and “peripheral” regions identified during recording. Six ROIs were selected from each region. Image analysis was performed using an ImageJ macro. Briefly, a nuclear count was performed by a particle count in the DAPI channel. A minimum intensity threshold was applied to the Alexa-568 channel, and this value was kept consistent within all ROIs from each tissue section. Activated caspase 3 staining was assessed by a particle count. Nuclear Ki67 staining was quantified by a particle count where the DAPI signal colocalised with the Alexa-568 signal. Particle counts were expressed as a percentage of the DAPI particle count in the same ROI to give a percentage of positively stained cells in each ROI for each antibody and averaged to give a single value for each section (one section per tumour).

### Measurement of intracellular pH

Cells were grown on glass coverslips for 48 h, then were incubated for 10 minutes at 21 °C in 1 μM 2′,7′-Bis(2-carboxyethyl)-5(6)-carboxyfluorescein acetoxymethyl ester (BCECF-AM, Biotium) in PSS, washed twice then left in PSS at pH 7.2 for 30 minutes before incubation for 10 minutes in PSS at pH 6.0 or 7.2. Coverslips were mounted in a Warner RC-20H recording chamber used in open configuration at room temperature with PSS perfusion at 1 ml/min. Two-point calibration was performed at the end of every experiment using K^+^-based PSS (where Na^+^ was replaced by K^+^) at pH 7.0 with 13 μM nigericin (Sigma) for 7 minutes, followed by K^+^-based PSS at pH 8.0 for a further 7 minutes. For each individual cell, a standard curve was derived using these buffers of known pH value, and the resting pH calibrated. Exposures of 0.15 s duration were taken every 15 s with a Nikon Eclipse TE200 epi-fluorescence microscope using SimplePCI 6.0 software to control the imaging system. Images were captured with a RoleraXR Fast1394 CCD camera (Q-imaging) with a 10X Plan Fluor objective. Images were saved as 16-bit .tif files and analysed in ImageJ 1.53c. Circular ROIs were placed over cells. The mean intensity at each wavelength was calculated for each ROI. Background fluorescence was calculated for each excitation wavelength by selecting a ROI containing no cells. Background fluorescence was subtracted from the mean intensity of each ROI before fluorescence ratio calculation. Each experimental repeat was the mean measurement from ∼40 cells per coverslip.

### Measurement of intracellular [Na^+^]

Cells were seeded at 2 x 10^4^ (MCF7) or 2.5 x 10^4^ (MDA-MB-231) cells/well in a 96 well, black walled, µclear polymer-bottomed plate (Greiner 655097). Medium was exchanged and drug incubations started after 36 h. Before dye loading, wells were washed with PBS, and 60 μl DMEM containing SBFI-AM (10 μM) and Pluronic F-127 (0.1%) ± drug treatment was added to each well. Cells were incubated in SBFI-AM at 37 °C for 2 h. Wells were then washed twice in PSS ± drug and left in PSS ± drug for imaging on a BMG Clariostar plate reader with excitation at 340 and 380 nm and emission collected at 510 nm. Background fluorescence was subtracted from each wavelength before fluorescence ratio calculation. Each experimental repeat was the mean fluorescence ratio of five wells from a single plate.

### Viability assay

Cells were cultured in 6 well plates. Culture medium from each well was removed from wells into 14 ml Falcon tubes then adherent cells were detached using trypsin-EDTA and added to the same tube. Cells were centrifuged at 800 x g for 5 minutes and resuspended in medium. A 10 μl sample was mixed with an equal volume of trypan blue and the number of viable and dead cells was counted using an Invitrogen Countess automated cell counter.

### Metabolic profiling

Oxygen consumption rate (OCR) and extracellular acidification rate (ECAR) measurements were conducted using a Seahorse XFe 96 Extracellular Flux Analyzer (Agilent), based on methods described previously (36, 37). The day prior to the measurements, MCF7 (2.0 x 10^4^/well) and MDA-MB-231 (3.0 x 10^4^/well) were seeded in a Seahorse XF96 cell culture microplate (101085-004, Seahorse) in 100 μl DMEM supplemented with 5% FBS and left at room temperature for 1 hour. Cells were then transferred to a 5% CO_2_ incubator set at 37 °C and incubated overnight. The next day, cells were washed twice with 200 μl of assay medium (DMEM powder, D5030, Sigma; resuspended in 1 L UF water, adjusted to pH 7.4, sterile-filtered and supplemented on the day of the experiment with 17.5 mM glucose, 2 mM glutamine and 0.5 mM sodium pyruvate) and a final volume of 180 μl was left per well. Then, the plate was incubated for equilibration in a 37°C non-CO_2_ incubator for 1 hour. In the meantime, the sensor cartridge was loaded with 20 μl of a 10X concentrated TTX solution to give a final assay concentration in the well of 30 μM. The experimental protocol consisted of consecutive cycles of 6 minutes that included 2 minutes of mixing followed by 4 minutes of measuring OCR and ECAR. After calibration of the cartridge and equilibration of the cell plate in the Seahorse Analyzer, basal measurements were acquired for 6 cycles, followed by the injection of the TTX or water vehicle control and measurement for 10 more cycles. There were 6 replicate wells per plate and the experiment was repeated independently 3 times.

### Human breast cancer tissue microarray

A tissue microarray consisting of formalin-fixed, paraffin-embedded cores from 1740 primary breast tumours was obtained from the BCNTB. Immunohistochemistry was performed as described previously (38). Samples were incubated with rabbit anti-human Na_v_1.5 antibody (1:100; Alomone ASC-013, which recognises residues 1978-2016 of human Na_V_1.5) and stained with EnVision+ Dual Link System/HRP (DAB+; Dako), following the manufacturer’s instructions and counterstained with Mayer’s haematoxylin. Specificity of staining was evaluated using antibody pre-adsorbed with immunising peptide. Slides were imaged on a Zeiss AxioScan.Z1 slide scanner with a 20X objective and staining was visualised using Zen 3.4 (blue edition) software (Zeiss). Staining was quantified using a modification to the Allred scoring system, as described previously (11, 39). TKL did the scoring and 10% of the series was independently verified by WF. In both cases, scoring was performed without prior knowledge of the associated clinical data. Concordance between investigators was assessed using Cohen’s weighted kappa (0.814; 95% CI 0.766-0.865; P < 0.001) and Intraclass Correlation Coefficient (0.954; 95% CI 0.938-0.966; P < 0.001), indicating excellent agreement between scorers.

### Statistical analysis

Linear regression was used for calibration of ratiometric indicators and pH electrodes over the recording ranges used. Statistical analysis was performed on raw (non-normalized) data using GraphPad Prism 9 for most analyses. IBM SPSS Statistics 27 was used to compute Intraclass Correlation Coefficient, Cohen’s weighted kappa and perform Cox multivariate proportional hazard analysis. Pairwise statistical significance was determined with Student’s paired or unpaired, or one-sample *t* tests for normally distributed data and Mann-Whitney tests for non-parametric data. Multiple comparisons were made using ANOVA and Tukey’s or Dunnett’s multiple comparisons tests for normally distributed data and using Kruskal-Wallis or Friedman’s tests for non-parametric data. Seahorse data were analysed using 2-way ANOVA. Time to event data were analysed using Kaplan Meier plots and log-rank (Mantel-Cox) tests computed with hazard ratio (HR) and 95% confidence intervals (CI). Correlations between Na_v_1.5 and other protein markers in the tissue microarray were evaluated using Spearman’s test. Results were considered significant at P < 0.05 or Benjamini-Hochberg (BH) adjusted P < 0.05 for differential gene expression.

## Results

### Na_v_1.5 protein expression associates with poor clinical outcome in breast cancer patients

Expression of Na_v_1.5 protein in breast cancer has previously been demonstrated in a small, qualitative study of 6 patients (12) and later in a study of 36 patients (11). To test the prognostic value of Na_v_1.5 in a larger cohort of patients, we used a breast cancer tissue microarray containing 1740 cases. Specificity of staining was confirmed by pre-incubation with the immunising peptide (Figure 1A). To explore correlation between Na_v_1.5 expression and histoclinical characteristics of the patient population, staining scores between 0-3 were classed as ‘low’ and scores between 4-8 were classed as ‘high’ (Figure 1A).

**Figure 1.**
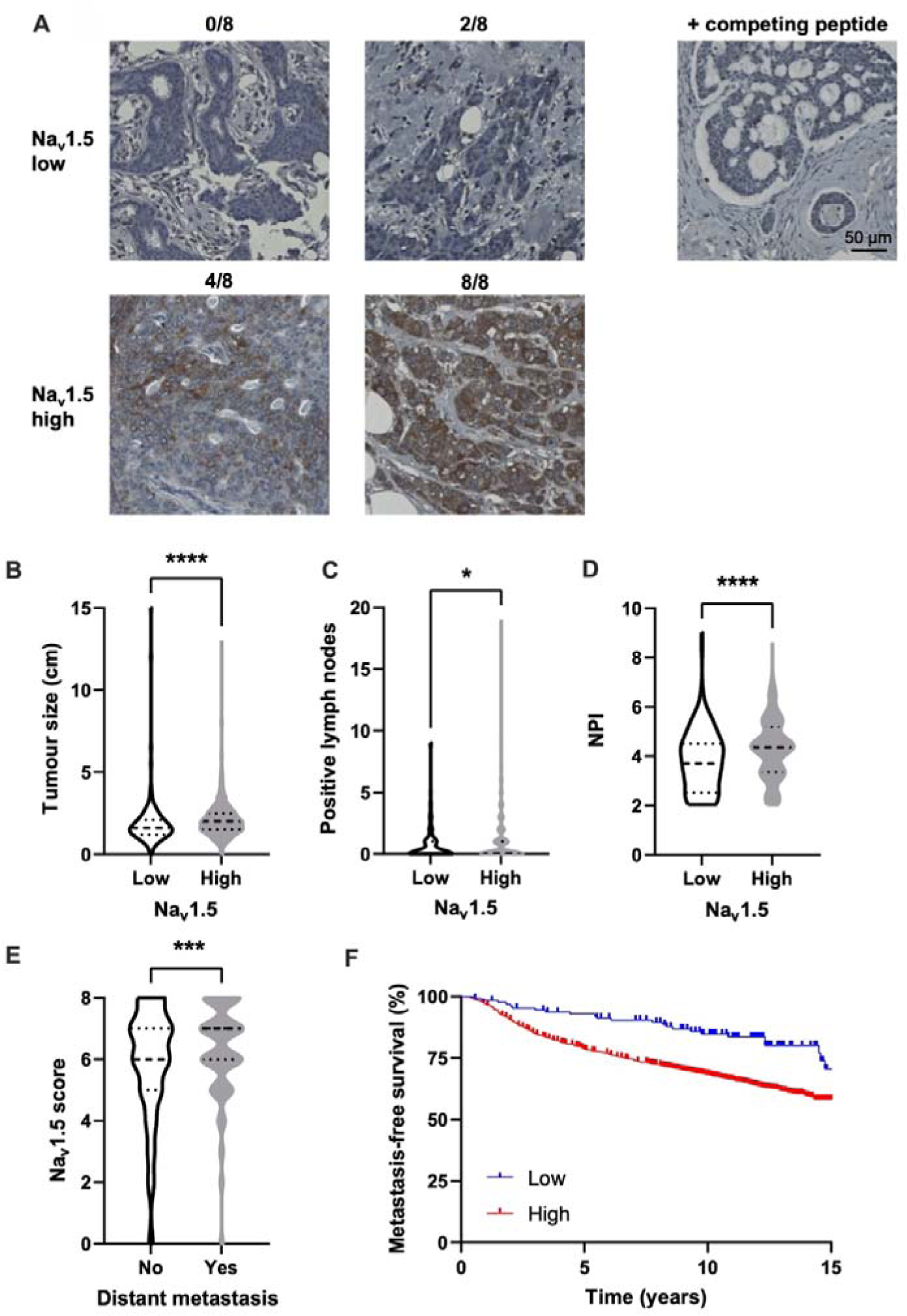
Na_v_1.5 expression in a breast cancer tissue microarray. (A) Examples of low and high Na_v_1.5 staining in carcinoma cells scored using a modified Allred system. Right: staining using anti-Na_v_1.5 antibody which had been preincubated with the immunising peptide. (B) Tumour size compared to Na_v_1.5 score. (C) Number of affected lymph nodes compared to Na_v_1.5 score. (D) Nottingham Prognostic Index (NPI) compared to Na_v_1.5 score. (E) Na_v_1.5 score in patients with or without recorded distant metastasis. Data are median + quartiles. *P < 0.05, ***P < 0.001, ****P < 0.0001; Mann-Whitney U tests (n = 1740). (F) Metastasis-free survival compared to Na_v_1.5 score. HR = 2.18 (95% CI 1.63-2.92); P < 0.001; log-rank test.

High Na_v_1.5 protein expression was correlated with larger tumour size (P < 0.001; Mann-Whitney U test; Figure 1B), lymph node positivity (P < 0.05; Mann-Whitney U test; Figure 1C), higher Nottingham Prognostic Index (P < 0.001; Mann-Whitney U test; Figure 1D), and higher tumour grade (P < 0.001; χ^2^ test; Table 1). In addition, Na_v_1.5 expression was significantly higher in patients who developed a distant metastasis (P < 0.001; Mann-Whitney U test; Figure 1E). Na_v_1.5 was negatively associated with estrogen receptor (ER; P < 0.05; Fisher’s exact test; Table 1) and progesterone receptor (PgR; P < 0.001; Fisher’s exact test; Table 1) expression, but positively associated with human epidermal growth factor receptor 2 (HER2; P < 0.01; Fisher’s exact test; Table 1). There was no association between Na_v_1.5 expression and triple negative breast cancer (TNBC; P = 0.12; Fisher’s exact test; Table 1), age (P = 0.52; Fisher’s exact test; Table 1), menopause (P = 0.46; Fisher’s exact test; Table 1) or endocrine therapy (P = 0.51; Fisher’s exact test; Table 1). However, high Na_v_1.5 expression was correlated with recorded chemotherapy use (P < 0.05; Fisher’s exact test; Table 1). These relationships are also displayed as violin plots in Supplementary Figure 1A-L.

**Table 1.**
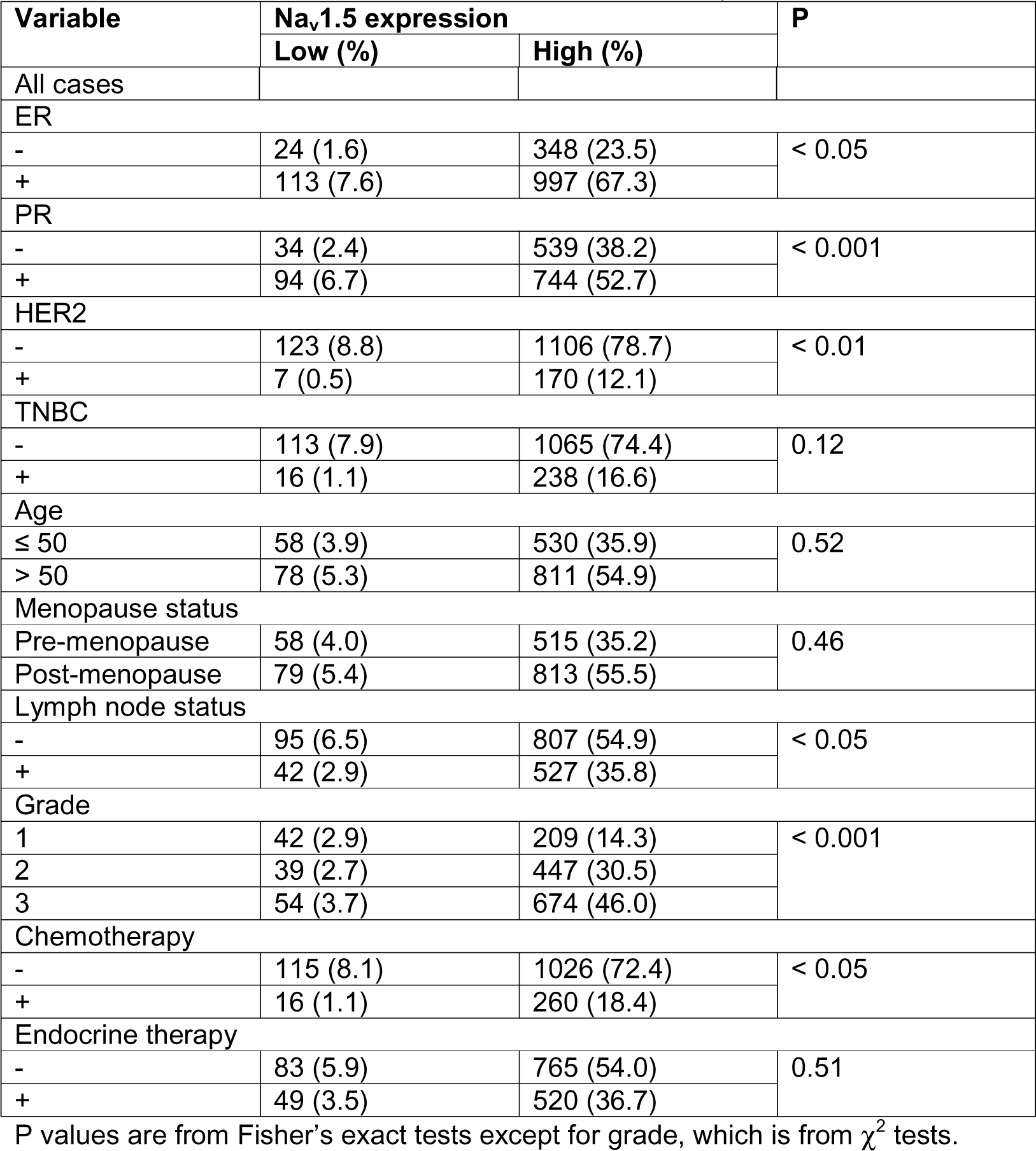
Patient histoclinical characteristics and Na_v_1.5 expression.

High Na_v_1.5 protein expression was associated with a significant reduction in metastasis-free survival (HR 2.18; 95% CI 1.63-2.92; P < 0.001; log-rank test; Figure 1F). This result was reflected in a significant reduction in overall survival (Supplementary Figure 2A), cancer-specific survival (Supplementary Figure 2B), disease-free survival (Supplementary Figure 2C), and local recurrence-free survival (Supplementary Figure 2D). When subdivided by receptor status, high Na_v_1.5 expression was associated with significantly reduced overall survival in ER+ (P < 0.05), but not HER2+ (P = 0.14) or TNBC patients (P = 0.53), although the sample sizes for HER2+ and TNBC patients were considerably smaller than for ER+ patients (Supplementary Figure 2E-G).

The prognostic value of Na_v_1.5 protein expression was considered in a Cox proportional hazards model including tumour size, grade, and lymph node status as categorical variables. Na_v_1.5 expression was an independent predictor of survival alongside the other variables in this model (HR = 1.58; 95% CI 1.05-2.37, P < 0.05; Table 2). Finally, the correlation between Na_v_1.5 expression and other protein markers previously scored in the same breast cancer tissue microarray was explored (40). This analysis revealed that Na_v_1.5 expression was significantly positively correlated with several other invasion-related protein markers (Table 3). In summary, high Na_v_1.5 protein expression associated with worse prognosis in combined subtypes of breast cancer patients across a range of clinical measures, highlighting the proposed role of this ion channel in promoting invasion and metastasis.

**Table 2.** Cox multivariate analysis of cancer-specific survival.

| Variable | Hazard ratio (95% CI) | P value |
| --- | --- | --- |
| Tumour size |  |  |
| ≤20 mm | 1 |  |
| >20 mm | 1.16 (1.10-1.22) | <0.001 |
| Grade |  |  |
| 1 | 1 |  |
| 2 | 1.46 (1.01-2.11) | <0.05 |
| 3 | 2.62 (1.86-3.70) | <0.001 |
| Lymph node status |  |  |
| Negative | 1 |  |
| Positive | 1.65 (1.36-2.00) | <0.001 |
| Na<sub>v</sub>1.5 expression |  |  |
| Low | 1 |  |
| High | 1.58 (1.05-2.37) | <0.05 |

**Table 3.** Correlation between Na_v_1.5 and other protein markers.

| **Protein marker** | **Spearman's correlation coefficient** | **P value** | **n** |
| --- | --- | --- | --- |
| ALDH1A1 | -0.05 | 0.265 | 579 |
| AMPK | -0.15 | < 0.001 | 1214 |
| Carbonic anhydrase IX | -0.07 | 0.111 | 479 |
| Cathepsin D | 0.11 | 0.003 | 762 |
| CD44 | -0.13 | 0.001 | 701 |
| CD56 | -0.19 | < 0.001 | 372 |
| HIF-1 | -0.05 | 0.576 | 130 |
| IDH2 | 0.13 | < 0.001 | 715 |
| Kank1M | 0.08 | 0.069 | 546 |
| Lactate dehydrogenase 5 | 0.00 | 0.942 | 434 |
| MMP11 | -0.02 | 0.849 | 117 |
| MMP13 (cytoplasmic) | -0.02 | 0.475 | 977 |
| MMP13 (nuclear) | -0.20 | < 0.001 | 977 |
| MMP14 (cytoplasmic) | -0.07 | 0.033 | 868 |
| MMP14 (membrane) | -0.09 | 0.009 | 867 |
| MMP14 (nuclear) | 0.01 | 0.754 | 867 |
| MMP15 (cytoplasmic) | -0.05 | 0.189 | 842 |
| MMP-15 (membrane) | 0.19 | < 0.001 | 842 |
| MMP15 (nuclear) | 0.19 | < 0.001 | 842 |
| MMP2 (cytoplasmic) | -0.20 | < 0.001 | 847 |
| MMP2 (nuclear) | -0.10 | 0.003 | 847 |
| MMP7 | -0.15 | < 0.001 | 866 |
| MMP9 | 0.11 | 0.007 | 599 |
| MX1 | 0.05 | 0.157 | 741 |
| NHERF1 | 0.07 | 0.214 | 321 |
| SNAT2 | -0.12 | 0.005 | 593 |

### Cells in patient breast cancer tissue exhibit voltage-sensitive inward and outward membrane currents

Much work has shown the potential for VGSCs to increase invasion and metastasis in cell culture models of breast and other epithelial cancers but until now, no electrophysiological recordings of VGSC currents have been shown in tissues taken directly from cancer patients. To address this, we performed electrophysiological experiments to record membrane currents in fresh tissue samples from three breast tumours (Supplementary Table 1). In fresh tissue slices made from these patient tumour biopsies, there were few cellular areas and most of the slices were composed of connective tissue or fat. Thus, we took patch clamp recordings from pockets of cells within the connective tissue at the top surface of the slice. Adipocytes were identified based on their large size and avoided. Portions of each tumour were also dissociated and seeded onto coverslips to enable patch clamp recording from isolated cells.

Voltage clamp recording revealed that cells in tumour slices expressed both voltage-sensitive inward and outward currents (Figure 2A). Small voltage-sensitive inward currents were found in two out of three patient specimens and in 4/17 recordings made (Figure 2B). The mean inward current-voltage relationship was noisy due to the small current density; however, it displayed activation at approximately −50 mV (Figure 2C), consistent with VGSC currents recorded from breast cancer cell lines (14, 41).

**Figure 2.**
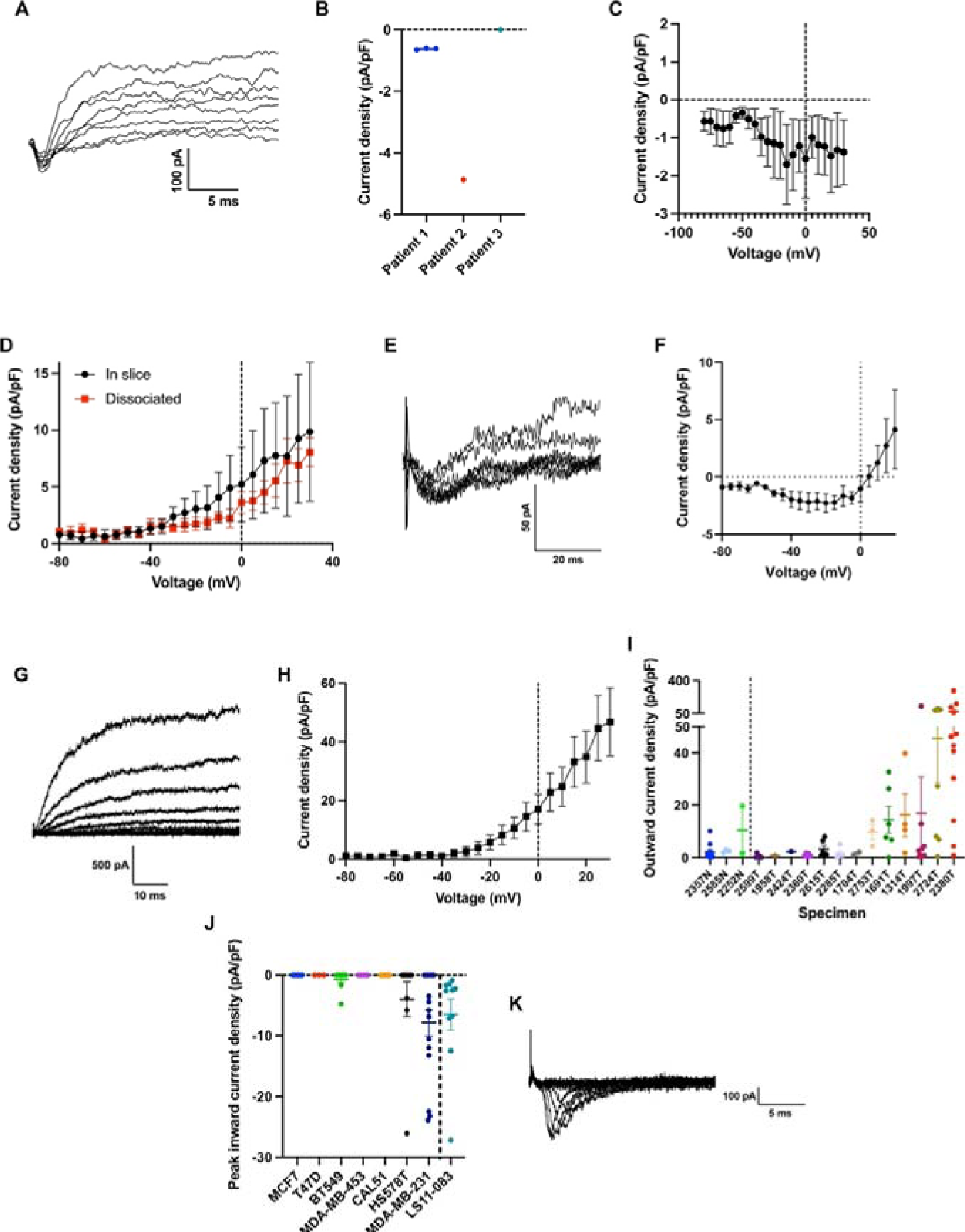
Characterisation of membrane currents in breast cancer tissue, dissociated primary cells, and cell lines. (A) Example inward and outward membrane currents recorded from a cell in a patient tumour slice, from a holding potential of −120 mV with 10 mV depolarising steps in the range −40 mV to +30 mV. (B) Peak inward current density, subdivided by patient tumour specimen. (C) Mean inward current-voltage relationship including cells from both patient tumour specimens (n = 4). (D) Mean outward current-voltage relationship from cells in patient tumour slices compared to cells which had been dissociated and plated onto glass coverslips (in slice: n = 10 cells from 3 patients; dissociated: n = 6 cells from 2 patients). (E) Example inward currents recorded from a dissociated mammary carcinoma cell, elicited by depolarising steps between −40 mV and −5 mV from a holding potential of - 120 mV. (F) Mean inward current-voltage relationship from dissociated mammary carcinoma cell samples (n = 4 cells). (G) Example outward currents recorded from a dissociated mammary carcinoma cell, using a holding potential of −120 mV with 10 mV depolarising steps in the range −80 mV to +30 mV. (H) Mean outward current-voltage relationship from dissociated mammary carcinoma cells (n = 30 cells). (I) Outward current density at +30 mV of dissociated normal mammary epithelial and carcinoma cells, subdivided by patient sample. (J) Peak inward current density measured across a panel of breast cancer cell lines and cancer-associated fibroblasts (CAFs). (K) Example voltage-sensitive inward currents in an LS11-083 CAF. Data are mean ± SEM.

Comparisons were then made between the cells in tumour slices, and cells dissociated from the same tumours plated onto glass coverslips. The mean outward current-voltage relationship was broadly similar between cells in slices and on coverslips (Figure 2D). It was not possible to further characterise the inward currents due to the small number of cells exhibiting these currents. In summary, both outward and inward voltage-sensitive currents were detected in cells in breast cancer tissue slices taken direct from patients, although making such recordings was technically challenging due to limited availability and the fibrous composition of the tissue. Since dissociation of tissue into isolated cells did not appear to affect the electrophysiological recordings, we moved onto recording from primary cell cultures.

### Primary breast epithelial and carcinoma cells exhibit voltage-sensitive inward and outward membrane currents

We recorded membrane currents from a total of four normal human mammary epithelial cell samples and thirteen breast cancer samples enriched for carcinoma cells (Supplementary Table 2). Voltage-sensitive inward currents were present in two out of 13 breast cancer samples and none of four normal human mammary epithelial cell samples (Figure 2E). The mean current-voltage relationship of the inward currents displayed activation at approximately −50 mV, again consistent with VGSC currents recorded from breast cancer cell lines (Figure 2F) (14, 41). Conversely, voltage-sensitive, non-inactivating outward currents were present in all normal human mammary epithelial cell samples tested and 10 out of 13 breast cancer samples (Figure 2G-I). In summary, (i) non-inactivating voltage-gated outward currents were common in normal human mammary epithelial cells and primary breast cancer cells whilst (ii) voltage-gated inward currents were rarely detectable, and only in malignant cells. The samples did not contain enough viable cells to allow further characterisation of the membrane currents or quantification of protein levels, however patch clamping is a more sensitive measure of plasma membrane ion channel expression than Western blot, given the rarity of these proteins.

### Inward currents are present in several triple-negative breast cancer cell lines and cancer-associated fibroblasts

To further explore the presence of VGSC currents in breast cancer cells, we set out to record from a panel of breast cancer cell lines and a breast tumour-derived cancer-associated fibroblast (CAF) cell line. Inward currents were present in MDA-MB-231 cells, consistent with previous reports (12, 14), and were also detected in Hs578T and BT549 cells (Figure 2J). Notably, all three cell lines in which inward currents were detected are from TNBC. In addition, inward currents were present in the CAF line LS11-083, with activation at around −25 mV (Figure 2K). Thus, we found functionally active VGSC currents in some TNBC and CAF cell lines, in broad agreement with several previous mRNA, protein, and electrophysiology studies (10–12, 14).

### Na_v_1.5 regulates expression of migration and invasion-promoting genes

In the tissue microarray study, we showed that higher Na_v_1.5 protein expression correlated with increased metastasis and more invasive tumours (Figure 1). In accordance with this, Na_v_1.5 activity has previously been shown to increase invasion and metastasis in preclinical models of breast cancer (11, 12, 14, 16–18, 24). Because of these findings, we hypothesised that knockdown of Na_v_1.5 would suppress expression of invasion-related genes. To test this hypothesis, we next compared gene expression in six MDA-MB-231 xenograft tumours with six tumours in which *SCN5A* expression had been stably suppressed using shRNA (11). We previously showed that Na_v_1.5 protein expression and Na^+^ current was ablated in these cells (11). Principal component analysis (PCA) showed that the *SCN5A* knockdown explained 25% of the variance and there was no relationship to mouse cage/block (Supplementary Figure 3A,B).

The *SCN5A* shRNA tumours displayed 136 differentially expressed genes, compared to the control tumours (BH-adjusted P < 0.05; Figure 3A). We next performed GSEA to evaluate the relationship between *SCN5A* knockdown and invasion-promoting genes (42). We used the MSigDB gene set SCHUETZ_BREAST_CANCER_DUCTAL_INVASIVE_UP which describes genes up-regulated in invasive ductal carcinoma vs ductal carcinoma in situ, a non-invasive type of breast tumour (43). GSEA of the differential expression in *SCN5A* shRNA tumours showed a significant reduction in invasion transcriptional response (P < 0.001; Figure 3B). There was also a smaller reduction in expression of invasion-downregulated genes (normalised enrichment score −2.35 vs. −3.58), although this was only significant at a reduced stringency (P < 0.05; Supplementary Figure 3C). These results support the notion that Na_v_1.5 is a key driver of invasion in breast cancer cells.

**Figure 3.**
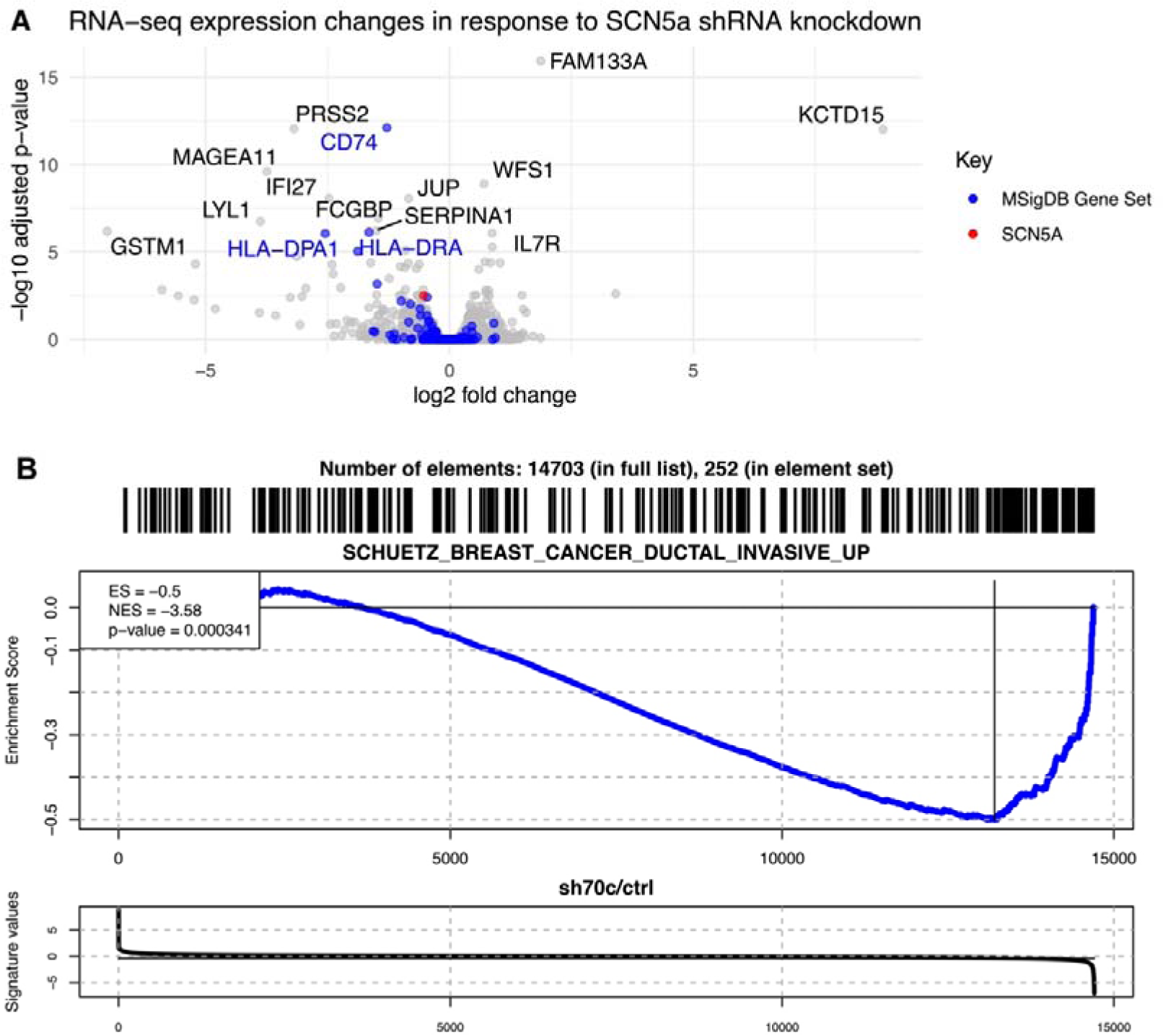
Altered human gene expression in Na_v_1.5 knockdown MDA-MB-231 xenograft tumours compared to control MDA-MB-231 tumours. (A) Volcano plot showing transcriptome changes in Na_v_1.5 knockdown tumours, with genes involved in invasion, and *SCN5A*, highlighted. (B) Gene set enrichment analysis (GSEA) of invasion-upregulated genes defined in the MSigDB gene set SCHUETZ_BREAST_CANCER_DUCTAL_INVASIVE_UP analysed in Na_v_1.5 knockdown vs. control tumours (n = 6 tumours in each condition).

### Na^+^ influx via Na_v_1.5 promotes glycolytic H^+^ production

A common feature of many types of cancer cell is a tendency to shift their metabolism to aerobic glycolysis and therefore H^+^ production (44). In MDA-MB-231 cells, Na_v_1.5 activity has been shown to increase H^+^ extrusion via NHE1, thus providing a lower pH_e_ optimal for cysteine cathepsin-dependent extracellular matrix degradation and increased invasion (23, 24, 45). However, the mechanistic link between Na_v_1.5 activity and NHE1-mediated H^+^ extrusion is unclear, given that Na^+^ influx through Na_v_1.5 would be expected to reduce the driving force for H^+^ export via NHE1. One possible explanation for the observed effect is that Na^+^ influx via Na_v_1.5 increases NKA activity to remove additional intracellular Na^+^ (46, 47) and maintain homeostasis. NKA has been shown to use ATP predominantly derived from glycolysis in many tissues including breast cancer cells (37, 48, 49). This dependence on glycolysis has been proposed to occur because it produces ATP close to the plasma membrane and the rate can quickly increase to cope with fluctuating demands from plasma membrane pumps (37). Thus, we hypothesised that Na^+^ entry through Na_v_1.5 would lead to an increase in glycolytic rate and acidic metabolite production. To test this hypothesis, we firstly examined the source of ATP used by NKA in breast cancer cells. As expected, the NKA inhibitor ouabain (300 nM; 6 h incubation) increased [Na^+^]_i_, measured using the ratiometric Na^+^ indicator SBFI-AM, by 2.2-fold in MDA-MB-231 cells (P < 0.001; n = 6; one-sample *t* test; Figure 4A) and by 2.0-fold in MCF7 cells (P < 0.001; n = 6; one sample *t* test; Figure 4B). This indicated that a rise in [Na^+^]_i_ could be used as a proxy for NKA inhibition. Interestingly, complete inhibition of mitochondrial respiration using the ATP synthase inhibitor oligomycin (1 µM; 2 h) did not alter [Na^+^]_i_ in MDA-MB-231 (P = 0.62; n = 6; one sample *t* test; Figure 4A) or MCF7 cells (P = 0.14; n = 6; one sample *t* test; Figure 4B). However, inhibition of glycolysis with the GAPDH inhibitor sodium iodoacetate (2 mM; 2 h) significantly increased [Na^+^]_i_ in both cell lines (P < 0.01 for both cell lines; n = 6; one sample *t* test; Figure 4A, B). These data suggest that NKA was able to maintain a steady [Na^+^]_i_ in the absence of mitochondrial respiration but not in the absence of glycolysis. Oligomycin had no effect on the viability of either cell line (P = 0.74 for MDA-MB-231 and P = 0.84 for MCF7; n = 3; one sample *t* tests), whereas sodium iodoacetate significantly reduced viability in both cell lines (P < 0.01 for MDA-MB-231 and P < 0.05 for MCF7; n = 3; one sample *t* tests; Supplementary Figure 4). Thus, these data suggest that in both MDA-MB-231 and MCF7 cells, NKA activity requires ATP derived from glycolysis to export Na^+^, and it can function without mitochondrial respiration.

**Figure 4.**
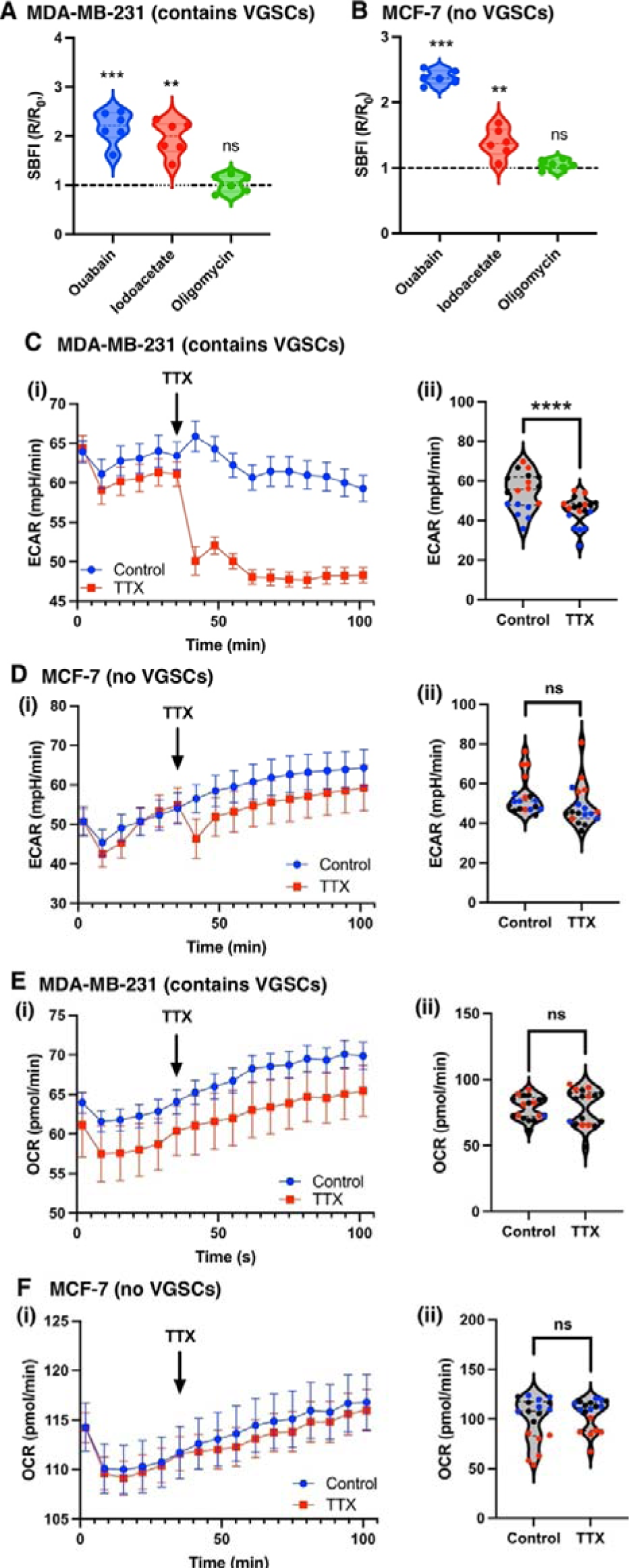
Effect of voltage-gated Na^+^ channel activity on glycolytic H^+^ production. (A) SBFI-AM fluorescence ratios of MDA-MB-231 cells after 6 h treatment with ouabain (300 nM), or 2 h treatment with sodium iodoacetate (2 mM) or oligomycin (1 µM). Data are normalised to the ratio in vehicle-treated cells (n = 6 experimental repeats; one sample *t* tests). (B) SBFI-AM fluorescence ratios of MCF7 cells after 6 h treatment with ouabain (300 nM), or 2 h treatment with sodium iodoacetate (2 mM) or oligomycin (1 µM). Data are normalised to the ratio in vehicle-treated cells (n = 6 experimental repeats; one sample *t* tests). (Ci) Representative measurements of the extracellular acidification rate (ECAR) of MDA-MB-231 cells. TTX was added to the treated cells at the indicated timepoint, to give a final concentration of 30 µM (n = 6 wells). (Cii) ECAR of MDA-MB-231 cells compared between control and TTX (30 µM) wells. (Di) Representative measurements of the ECAR of MCF7 cells. TTX was added to the treated cells at the indicated timepoint, to give a final concentration of 30 µM (n = 6 wells). (Dii) ECAR of MCF7 cells compared between control and TTX (30 µM) wells. (Ei) Representative measurements of the oxygen consumption rate (OCR) of MDA-MB-231 cells. TTX was added to the treated cells at the indicated timepoint, to give a final concentration of 30 µM (n = 6 wells). (Eii) OCR of MDA-MB-231 cells compared between control and TTX (30 µM) wells. (Fi) Representative measurements of the OCR of MCF7 cells. TTX was added to the treated cells at the indicated timepoint, to give a final concentration of 30 µM (n = 6 wells). (Fii) OCR of MCF7 cells compared between control and TTX (30 µM) wells. Each data point in panels (ii) represents the mean of the last 6 timepoints for each well (n = 3 experimental repeats each containing 6 wells; experimental repeats colour-coded black, red, blue). Data are mean ± SEM. ****P < 0.0001, ***P < 0.001, **P < 0.01, *P < 0.05, ns, not significant (2-way ANOVA).

Next, we used a Seahorse XFe96 analyser to test the hypothesis that Na^+^ influx through Na_v_1.5 increases the rate of glycolysis via increasing ATP demand from NKA. The effects of the VGSC inhibitor TTX (30 µM) on the extracellular acidification rate (ECAR; a measure of glycolysis) and oxygen consumption rate (OCR; a measure of mitochondrial respiration) were compared between MDA-MB-231 cells (which express functional Na_v_1.5 channels) and MCF7 cells (which do not display these currents; Figure 2J). In control experiments, addition of TTX to wells containing medium without cells transiently reduced the measured ECAR, which then returned to baseline levels within 15 minutes (Supplementary Figure 5). Therefore, measurements on cells ± TTX were compared after this period. Addition of TTX (30 µM) to MDA-MB-231 cells caused a rapid and sustained reduction in ECAR (P < 0.0001; n = 3 experimental repeats containing 6 wells each; 2-way ANOVA; Figure 4C). However, TTX had no significant effect on the ECAR of MCF7 cells, which do not express functional Na_v_1.5 channels (50) (P = 0.07; n = 3 experimental repeats containing 6 wells each; 2-way ANOVA; Figure 4D). In contrast, OCR was unaffected by TTX in both cell lines (P = 0.99 and 0.11 for MDA-MB-231 and MCF7, respectively; n = 3 experimental repeats containing 6 wells each; 2-way ANOVA; Figure 4E,F). Together, these data are consistent with Na^+^ influx via Na_v_1.5 increasing the rate of glycolysis but not mitochondrial respiration. This result can therefore explain the established link between Na_v_1.5 activity and NHE1-induced extracellular acidification which promotes invasion. It also highlights a novel link between Na^+^ homeostasis and altered metabolism in cancer cells.

### Extracellular pH is lower towards the periphery of xenograft tumours and low pH correlates with high cellularity and proliferation

The tumour microenvironment is reported to be acidic (51), with a low intratumoural pH facilitating various metastatic hallmarks including ECM degradation and invasion. Classically, this low pH has been thought to be due to hypoxia in the poorly perfused areas of the tumour. Here we show evidence that low extracellular pH can instead be associated with highly proliferative, peripheral areas of the tumour where it would be expected that there is increased metabolic activity. This would be consistent with areas of high glycolytic activity, and potentially high NKA activity. We therefore next assessed the extracellular pH (pH_e_) of MDA-MB-231 xenograft tumours using pH-sensitive microelectrodes. Measurements were recorded in several locations on the top surface of tissue slices directly prepared from xenograft tumours, alternately from the opaque core of the tumour, and from more translucent periphery (Figure 5A). The mean overall pH_e_ was 6.9 ± 0.1, which is significantly lower than pH 7.4 normally found in the extracellular fluid of healthy tissue (P = 0.001; n = 9; one sample *t* test) (51). The mean pH_e_ in the core was 7.0 ± 0.1; in the periphery, it was significantly lower, at 6.8 ± 0.1 (P < 0.01; n = 9; paired *t* test; Figure 5B). The differences between core and periphery were investigated in more detail using immunohistochemistry (Figure 5C). Cellularity (mean nuclear count per ROI) was significantly higher in the periphery compared to the core (988 ± 29 vs. 838 ± 47; P < 0.01; n = 9 tumours; paired *t* test; Figure 5D). Similarly, proliferation, as measured by Ki67-positive nuclei, was significantly higher in the periphery than the core (20.1 ± 7.0% in the periphery vs. 6.9 ± 2.4% in core; P < 0.05; n = 9 tumours; paired *t* test; Figure 5E). Conversely, apoptosis, as measured by cleaved caspase 3 positivity, was significantly higher in the core compared to the periphery (10.9 ± 3.5% in core vs. 2.1 ± 0.7% in periphery; P < 0.05; n = 9 tumours; paired *t* test, Figure 5F). Taken together, these results suggest that in this model the pH_e_ is lower in peripheral regions with high cellularity which are proliferating rapidly, and higher towards the core of the tumours, where there is more apoptosis.

**Figure 5.**
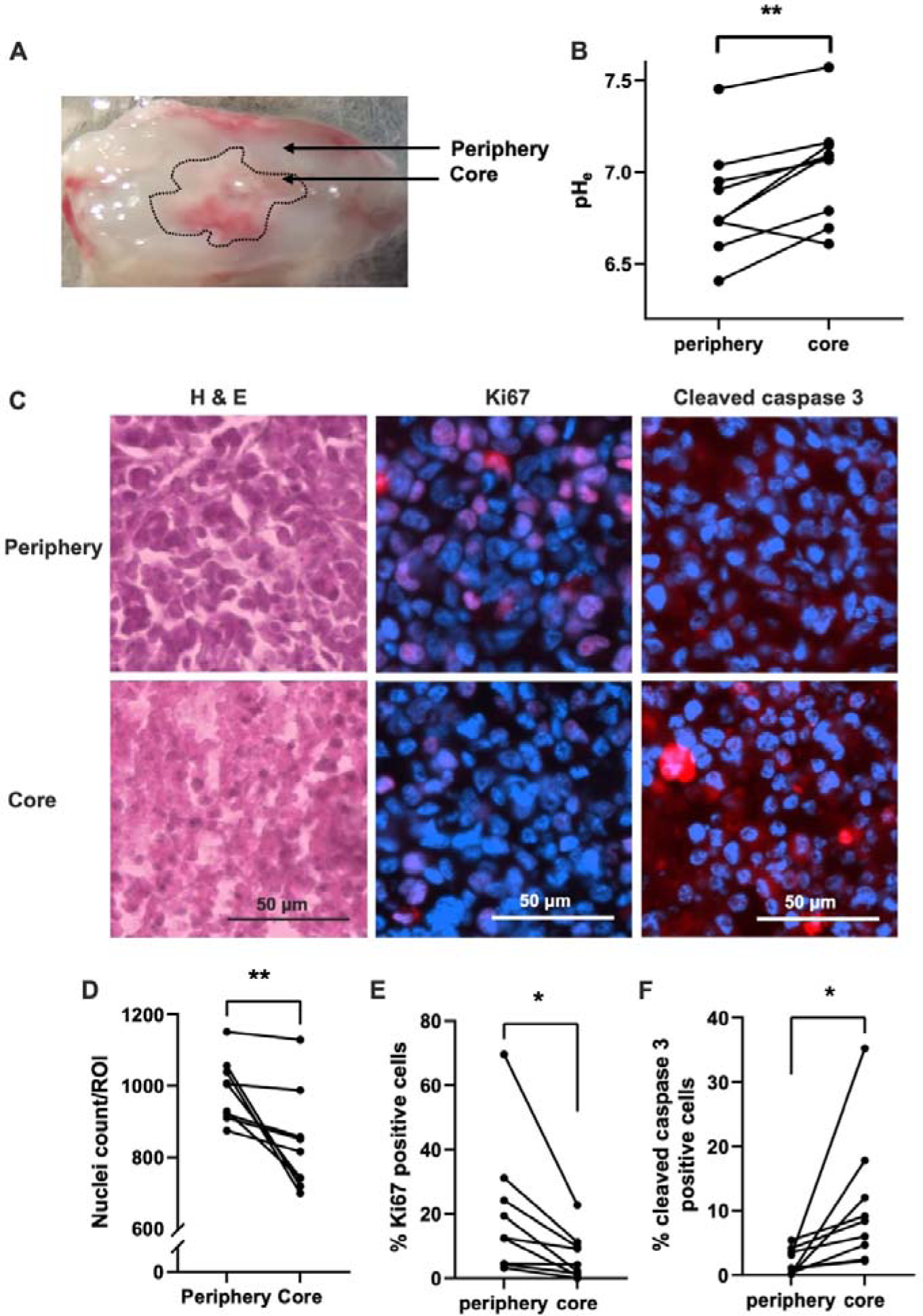
Altered extracellular pH in tissue slices from MDA-MB-231 xenograft tumours. (A) Photograph of a tumour slice showing the difference in appearance between the translucent periphery and opaque core. (B) Tumour slice extracellular pH (pH_e_) comparing core and peripheral regions in each slice; n = 9 tumours (one slice from each). (C) Examples of peripheral and core regions with H&E staining (left), Ki67 (red, middle), and cleaved caspase 3 (red, right) with DAPI (blue). (D) Cellularity in the peripheral and core regions of each section, based on DAPI nuclear count (n = 9 tumours). (E) Ki67-positive cells (%) in peripheral and core regions (n = 9 tumours). (F) Cleaved caspase 3-positive cells (%) in peripheral and core regions (n = 9 tumours). Data are mean ± SEM; *P < 0.05, **P < 0.01 (paired *t* tests).

### Persistent Na_v_1.5 current is increased by low extracellular pH

Given that the tumour pH_e_ is acidic *in vivo*, we next assessed the effect of pH alteration on Na_v_1.5 activity in MDA-MB-231 cells using patch clamp recording. A pH_e_ of 7.2 was compared with a pH_e_ of 6.2 since the pH_e_ of solid tumours has been reported to approach this level of acidity (52). Lowering pH_e_ to 6.2 reduced the transient Na^+^ current from −13.5 ± 2.3 pA/pF to −9.6 ± 1.5 pA/pF (P < 0.01; n = 11; Wilcoxon matched pairs test; Figure 6A, C). In contrast, the persistent Na^+^ current, measured at 20-25 ms after depolarisation, was increased by acidification to pH_e_ 6.2, from −0.31 ± 0.04 pA/pF to −0.71 ± 0.11 pA/pF (P < 0.01; n = 10; paired *t* test; Figure 6B, D). Analysis of the voltage dependence of activation and steady-state inactivation (Figure 6E-G) revealed that the slope factor (k) and voltage at half activation (V½) did not significantly change when the pH_e_ was reduced from 7.2 to 6.2 (P = 0.077 and 0.087, respectively; n = 10; paired *t* tests, Table 4). However, the V½ was significantly depolarised at pH_e_ 6.2, from −80.4 ± 1.4 mV to −73.3 ± 2.8 mV (P < 0.01; n = 10; paired *t* test; Table 4) and the k for inactivation was also significantly changed, from −8.4 ± 0.8 mV to −11.9 ± 0.9 mV (P < 0.01; n = 10; paired *t* test; Table 4). This depolarising shift in steady-state inactivation thus increased the size of the window current (Figure 6G, H). Indeed, at the reported resting V_m_ of MDA-MB-231 cells, −18.9 mV (12), reducing the pH_e_ from 7.2 to 6.2 more than doubled channel availability from 1.9 ± 0.6% to 4.9 ± 0.7% of maximum (P < 0.05; n = 10; paired *t* test; Figure 6I). When pH_e_ was reduced further to 6.0, the effect on channel availability was even greater, increasing nearly five-fold from 2.1 ± 0.9% at pH_e_ 7.2 to 10.3 ± 2.2% of maximum at pH_e_ 6.0 (P < 0.001, n = 8, paired *t* test; Figure 6J & Supplementary Figure 6A, B).

**Figure 6.**
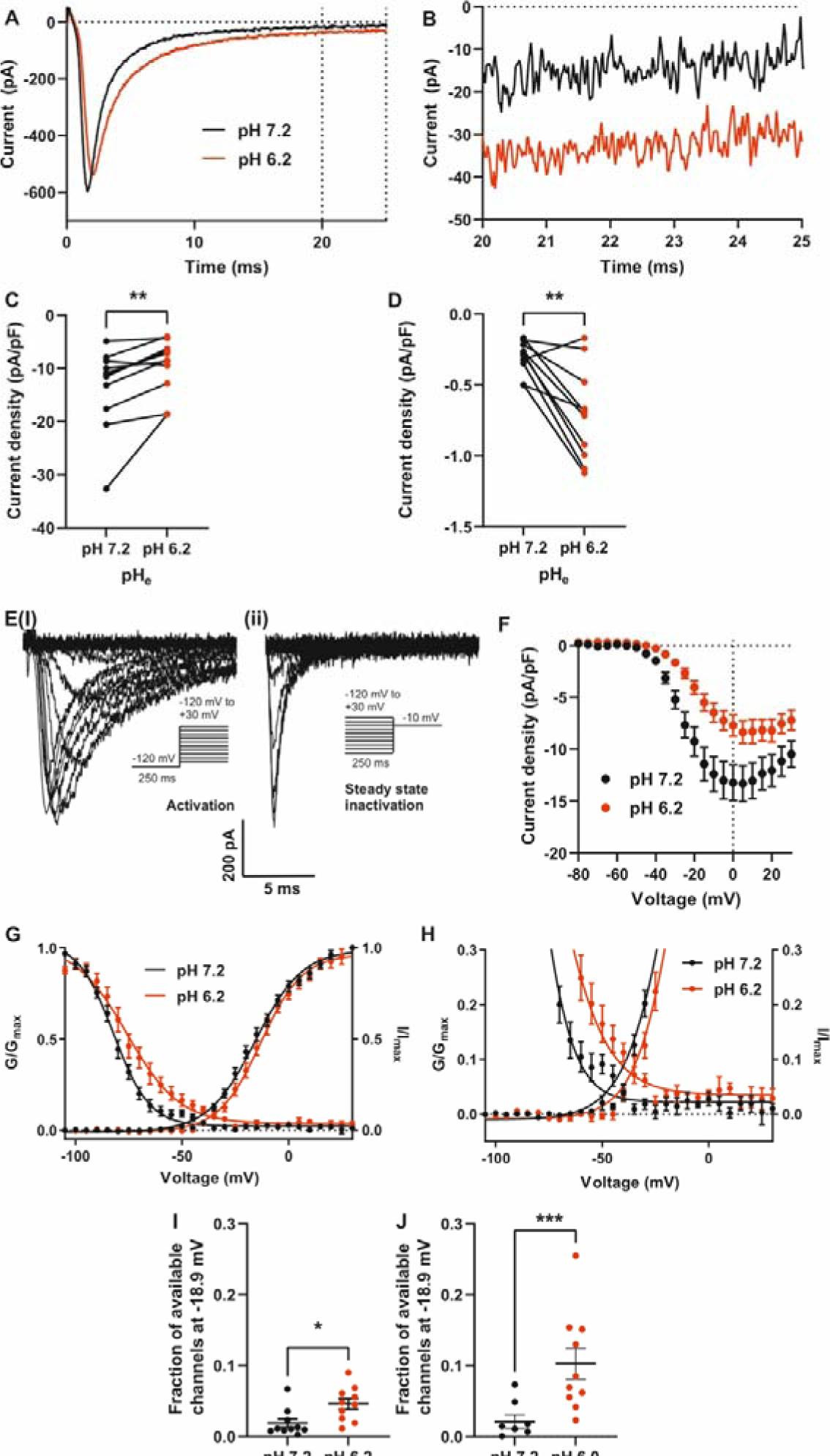
Effect of low pH_e_ on Na^+^ current in MDA-MB-231 cells. (A) Example Na^+^ currents elicited by depolarisation to 0 mV from a holding voltage of −120 mV. (B) Data from (A) expanded between 20 and 25 ms after depolarisation. (C) Peak Na^+^ current density (P < 0.01; n = 11; Wilcoxon matched pairs test). (D) Mean persistent Na^+^ current density measured between 20 and 25 ms after depolarisation (P < 0.01; n = 10; paired *t* test). (E)(i) Example family of Na^+^ currents generated by the activation voltage clamp protocol (inset). (ii) Example family of Na^+^ currents generated by the steady-state inactivation voltage clamp protocol (inset). (F) Current density-voltage relationship (n = 17). (G) Overlay of activation and inactivation curves at pH 7.2 and 6.2 (n = 10 cells with the largest currents). (H) Expanded data from (G) showing the window current. (I) Fraction of channels available at the reported resting membrane potential of −18.9 mV at pH 7.2 and 6.2 (n = 10 cells with the largest currents; *t* test). (J) Fraction of channels available at the reported resting membrane potential of −18.9 mV, at pH 7.2 and 6.0 (P < 0.001; n = 8 cells; *t* test). Data are mean ± SEM.

**Table 4.** Effect of reduced pH_e_ on VGSC Na^+^ current parameters.

| Parameter | pH 7.2 | pH 6.2 | P | n |
| --- | --- | --- | --- | --- |
| Peak current density (pA/pF) | -13.5 ± 2.3 | -9.6 ± 1.5 | 0.006 | 11 |
| Persistent current density (pA/pF) | -0.31 ± 0.04 | -0.71 ± 0.11 | 0.006 | 10 |
| Activation V<sub>1/2</sub> (mV) | -15.2 ± 2.0 | -12.9 ± 1.8 | 0.087 | 10 |
| Activation k (mV) | 10.7 ± 0.5 | 9.5 ± 0.4 | 0.077 | 10 |
| Inactivation V<sub>1/2</sub> (mV) | -80.4 ± 1.4 | -73.3 ± 2.8 | 0.003 | 10 |
| Inactivation k (mV) | -8.4 ± 0.8 | -11.9 ± 0.9 | 0.005 | 10 |

V<sub>1/2</sub>: half (in)activation voltage; k: slope factor for (in)activation. The holding potential was -120 mV. Results are mean ± SEM. Statistical comparisons were made with paired *t* tests on non-normalised data.

Cancer cells have been shown to have an inverted pH ratio across the plasma membrane, so the exterior of the cell is more acidic but the intracellular fluid has a higher pH than normal cells (51). For this reason, it was important to explore whether the pH_e_-dependent electrophysiological changes were mediated by a change in intracellular pH (pH_i_). To do this we assessed the effect of pH_e_ on pH_i_ using the ratiometric fluorescent pH indicator BCECF-AM to measure pH_i_ following incubation in different pH_e_. Lowering pH_e_ from 7.2 to 6.0 led to intracellular acidification to pH_i_ 6.3 ± 0.1 (Supplementary Figure 7A, B). However, altering pH_i_ from 7.2 to 7.6 (empirically the maximum range of intracellular patch pipette solution pH which still allowed the formation of giga-Ohm seals onto MDA-MB-231 cells) had no effect on transient or persistent Na^+^ current or voltage dependence of activation or steady-state inactivation (Supplementary Figure 7C-G). In summary, Na^+^ entry into breast cancer cells through Na_v_1.5 is increased in acidic pH_e_ but is not sensitive to changes in pH_i_ under the range of pH_i_ tested. These data suggest that areas of the tumour with lower pH_e_ would have increased persistent Na^+^ current into breast cancer cells at steady state. This additional Na^+^ influx would either lead to a slow and continuous increase in intracellular [Na^+^], resulting in cell death (the opposite of what we observed in the more acidic parts of the tumour), or it would lead to increased NKA activity in the more acidic parts of the tumour, to maintain a stable intracellular [Na^+^] and maintain cell viability.

### Prediction of Na_v_1.5-dependent extracellular acidification rate

The expected ECAR due to VGSC activity can be calculated if it is assumed that the persistent Na^+^ current into cells through VGSCs is counteracted by activity of NKA to maintain a stable [Na^+^]_i_. The other assumptions used in this calculation are that NKA pumps 3 Na^+^ ions out of the cell per cycle in which it hydrolyses one molecule of ATP (53). Glycolytic production of lactic acid produces 2 molecules of ATP and 2 molecules of lactate per glucose. The production of H^+^ by this reaction coupled to the ATPase hydrolysis of ATP to ADP generates 2 H^+^ per glucose (54). There is therefore net production of one H^+^ per ATP molecule generated by glycolytic fermentation to produce lactate. Using these assumptions, we calculated the ECAR due to Na_v_1.5 activity to be 1.3 mpH/min (full calculations delineated in Supplementary Materials). This predicted ECAR is within an order of magnitude of the measured change in ECAR due to TTX inhibition of Na_v_1.5 (9.8 ± 1.7 mpH/min; Figure 4D). Thus, our model can explain how Na_v_1.5 activity can increase H^+^ extrusion through NHE1, considering experimental variability in determination of ECAR and persistent Na^+^ current, and estimation of pH_e_ during the Seahorse assay.

### Protein-protein interactions of NHE1

NHE1 is the pH regulator which has been implicated as most important in Na_v_1.5-dependent extracellular acidification and protein-protein interactions have been suggested to play a significant role in regulating NHE1 activity, including by Na_v_1.5 (23, 55, 56). We therefore looked for other protein interactions of NHE1. Using the STRING database (v11.5) we searched for the top 50 likely protein interactions of NHE1, only considering the proteins sharing a physical complex. This identified several subunits of NKA as likely binding partners of NHE1 (Supplementary Figure 8). An interaction between NKA and NHE1 further supports a model in which NKA is an intermediate step by which Na^+^ influx through channels such as Na_v_1.5 can then alter NHE1 activity.

In summary, as well as identifying a mechanism by which Na_v_1.5 may increase tumour acidification via enhanced rate of glycolysis, our data suggest that the acidic tumour microenvironment could increase Na^+^ influx via Na_v_1.5. Together, these findings suggest that there is a positive feedback loop in breast cancer cells, whereby Na^+^ influx and H^+^ release into the tumour microenvironment could synergise to promote invasion and metastasis (Figure 7).

**Figure 7.**
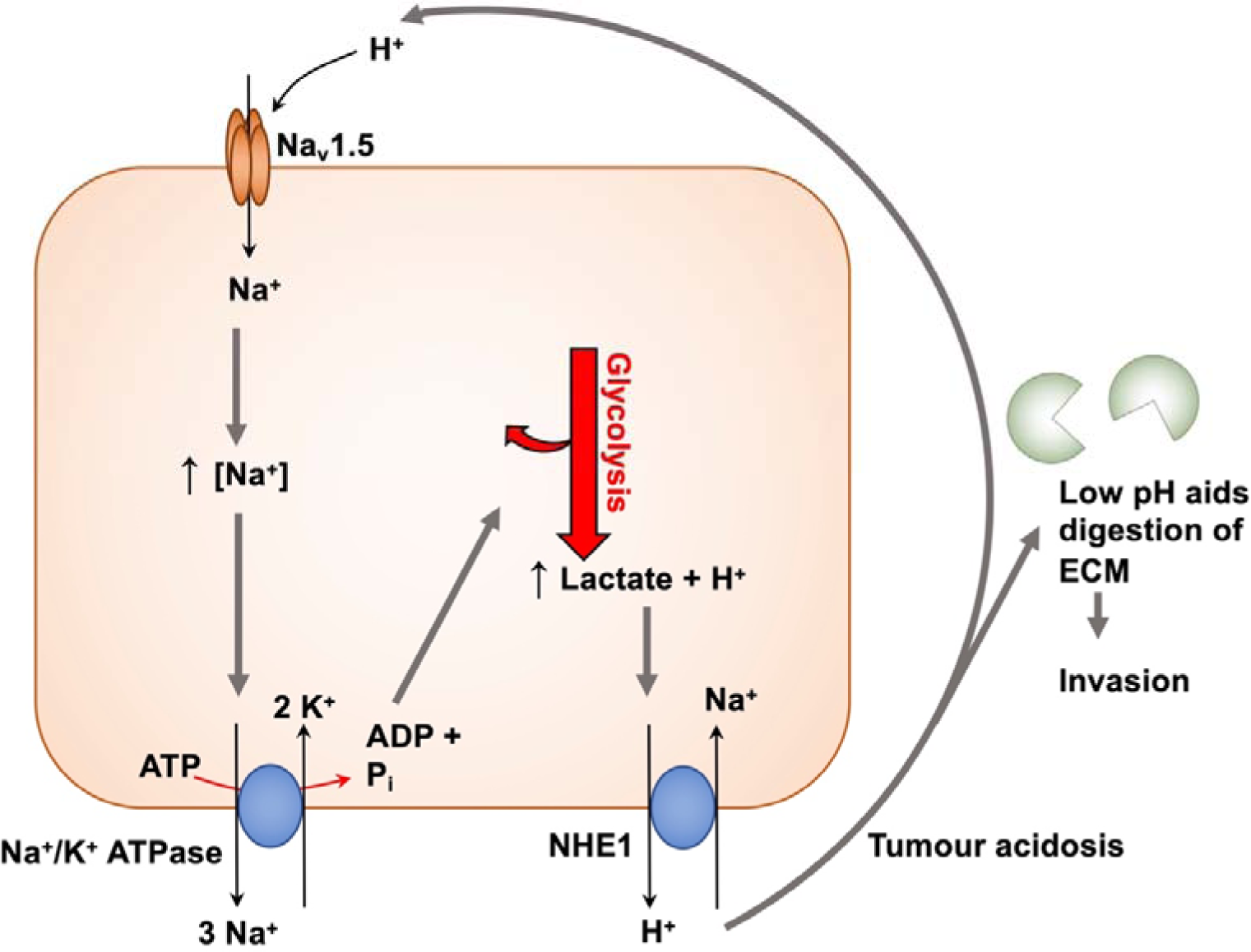
Mechanism for Na_v_1.5-mediated cellular invasion. Elevated steady-state Na^+^ entry through Na_v_1.5, leads to increased NKA activity. The increased ATP demand is satisfied by glycolysis, increasing H^+^ production and extrusion through pH regulators such as CAIX and NHE1. Extracellular acidification in turn increases persistent Na^+^ entry via Na_v_1.5, leading to a positive feedback loop linking Na_v_1.5, Na^+^ entry, increased H^+^ production via glycolysis, and extracellular acidification. This results in ECM degradation and increased invasion (7, 23, 45).

## Discussion

In this study on breast cancer, we show that upregulation of Na_v_1.5 protein expression is positively associated with metastasis and reduced cancer-specific survival. We also identify a novel activity-dependent positive feedback role for this channel which results in increased metabolic activity and extracellular acidification. We have thus provided an integrated mechanism by which Na_v_1.5 can promote metastatic dissemination. We show specifically that acidity of the tumour microenvironment, particularly in the invasive periphery of the tumour, enhances persistent Na^+^ current through Na_v_1.5 into breast cancer cells, and this in turn promotes glycolytic metabolic activity. We therefore propose that Na_v_1.5 is a key regulator of the ionic tumour microenvironment, facilitating local invasion during the early stage of metastasis.

### Clinical significance of Na_v_1.5 expression in breast cancer

Our findings support, for the first time, a pro-metastatic role for Na_v_1.5 in the clinical setting. We show that Na_v_1.5 protein expression correlates with lymph node positivity, increased metastasis, higher tumour grade and consequently reduced survival. In addition, the prognostic potential of Na_v_1.5 expression is independent of, but of comparable importance to, lymph node status, tumour grade and tumour size. Although there have been several previous reports indicating that VGSCs are likely to be important prognostic indicators of breast cancer (10–12), no study has previously examined Na_v_1.5 protein expression in a large cohort of breast cancer patients. Previously, VGSC protein expression in breast cancer has mostly been extrapolated from experiments in cell lines with differing metastatic potential (12, 14) and small cohorts of patients (11–13). Interestingly, we found that Na_v_1.5 expression was negatively associated with ER status, and positively correlated with HER2 status, but not with TNBC status. Together, these findings suggest possible functional linkage between ER, HER2 and Na_v_1.5 expression. In agreement with this notion, ER+ MCF-7 cells have low VGSC expression (12), and silencing ER in MCF-7 cells increases Na^+^ current and VGSC-dependent invasion (57). In addition, various tyrosine kinase receptors closely related to HER2, including EGFR, have been shown to regulate VGSC expression in carcinoma cells (58). Further work is required to determine how these receptors interact with Na_v_1.5 in breast cancer cells.

Na_v_1.5 protein expression does not necessarily result in active channels at the plasma membrane. Therefore, for the first time, we attempted to investigate functional channel activity at the plasma membrane using whole cell patch clamp recording. Although non-inactivating outward currents were widespread, small inward currents, indicative of Na_v_1.5 activity, were rarer and harder to detect. This result is surprising given the high proportion of Na_v_1.5 positive cells in the tissue microarray. This apparent contradiction may be explained if a large proportion of the channels are present on intracellular membranes. Another possibility is that the tissue slice, dissociation, and cell culture conditions resulted in transport of channels away from the plasma membrane, as has been shown for K_Ca_3.1 (59). Indeed, the cancerous tissues and primary cultures contained very few cells viable enough for electrophysiological recording. A few studies have reported the presence of Na^+^ currents carried by VGSCs in cells dissociated from mesothelioma and cervical tumour tissue, although the latter had been maintained in long-term culture (60–62). A recent study also demonstrated Na^+^ currents in primary colorectal carcinoma cells (55). To our knowledge, ours is the first report of VGSC currents in breast cancer tissue or primary cells from patients, however more work is required to refine the procedures for electrophysiological recordings using such clinical material.

### Na_v_1.5-induced extracellular acidification – a positive feedback mechanism promoting invasion

Our unexpected finding that the proliferating and invasive tumour periphery was more acidic than the hypoxic core agrees with a number of other studies (51, 63–66). Low extracellular pH in tumours promotes invasion by increasing activity of low pH-dependent enzymes that degrade the ECM, such as cysteine cathepsins (24, 45), so it makes sense that the invading edge of a tumour should have a low pH_e_. We found that low pH_e_, as found in tumours, increases the persistent Na^+^ current through Na_v_1.5 in breast cancer cells, thereby promoting Na^+^ influx. Greater Na^+^ influx into cancer cells in tumour regions of lower pH_e_ may be partially responsible for the heterogeneity of apparent tumour [Na^+^] as measured by ^23^Na-MRI (67, 68).

Increased Na_v_1.5-mediated Na^+^ influx into cancer cells would be expected to promote activity of NKA (69). There is substantial evidence that NKA utilises glycolysis as its main ATP source (21, 37, 70, 71). NKA activity, and consequent glycolytic metabolism, would therefore increase the rate of H^+^ production. In agreement with this paradigm, we found that Na_v_1.5 activity in breast cancer cells increased glycolysis, as measured by extracellular H^+^ production, without affecting oxidative phosphorylation. Increasing [Na^+^]_i_, via the ionophore gramicidin, has previously been shown to potentiate the rate of glycolysis in breast cancer cells, whereas the NKA inhibitor ouabain decreased H^+^ production (37). These findings echo those where glycolysis was found to be the ATP source for another plasma membrane ion pump, the plasma membrane Ca^2+^ ATPase (PMCA) in pancreatic cancer cells (72, 73). We found that inhibition of glycolysis induced a large increase in [Na^+^]_i_ similar to that caused by ouabain, and also rapidly led to cell death.

Elevated steady-state Na^+^ entry via persistent current through Na_v_1.5, leading to increased glycolysis to power NKA, would in turn, be expected to increase H^+^ extrusion through pH regulators such as CAIX and NHE1. Reciprocally, the reduction in pH_e_ serves to increase persistent Na^+^ entry into breast cancer cells via Na_v_1.5. Together, these mechanisms would lead to a positive feedback loop linking Na_v_1.5, Na^+^ entry, increased H^+^ production via glycolysis and extracellular acidification (Figure 7). This model fits with previous studies showing that Na_v_1.5 activity increases NHE1-mediated H^+^ extrusion in breast cancer cells, leading to ECM degradation and increased invasion (23, 45). It would also explain how Na^+^ influx via Na_v_1.5 increases H^+^ efflux via NHE1, despite an apparently wasteful collapse of the inward Na^+^ gradient that powers NHE1-mediated extrusion of H^+^ (7). Given our evidence that extracellular acidification in breast tumours occurs particularly in the highly proliferative peripheral region, and that the persistent Na^+^ current through Na_v_1.5 is larger in acidic conditions, more Na^+^ would be likely to enter breast cancer cells at the invading edges of the tumour.

## Conclusion

Here, we have shown that Na_v_1.5 is associated with poor prognosis and increased metastasis in breast cancer. Since Na_v_1.5 is a negative prognostic indicator, and its expression increases tumour growth and metastasis in preclinical models (11), it is a promising target for drug repurposing and discovery (10, 74, 75). In agreement with this notion, we recently showed that exposure to certain persistent Na^+^ current-inhibiting Class 1c and 1d antiarrhythmic drugs is associated with significantly improved cancer-specific survival (76). In addition, VGSC-inhibiting drugs have been shown to decrease tumour growth and metastasis in murine breast cancer models (17, 18). Furthermore, a recent clinical trial has shown that presurgical peritumoral treatment with lidocaine significantly improves disease-free and overall survival in women with early breast cancer (19). In conclusion, our results reveal a positive feedback mechanism by which Na^+^ influx through Na_v_1.5 promotes glycolytic H^+^ production to increase invasive capacity and drive breast cancer metastasis. This novel mechanism, together with the emerging clinical data, underscores the value of Na_v_1.5 as a prognostic marker and potential anti-metastatic therapeutic target.

## Acknowledgements

The authors wish to acknowledge the roles of the Breast Cancer Now Tissue Bank in collecting and making available the samples and data, and the patients who have generously donated their tissues and shared their data to be used in the generation of this publication. The authors also thank Prof Miles Whittington (Hull-York Medical School, UK), Dr John Davey and Dr Katherine Newling (Technology Facility, University of York, UK) and Prof Lýdia Vargová (Charles University, Czechia) for providing invaluable advice. For the purpose of open access, a Creative Commons Attribution (CC BY) licence is applied to any Author Accepted Manuscript version arising from this submission.

## Author contributions

Study conception and design: TKL, SC, WJB. Data collection: TKL, AT, ADJ, SPF, NS, MAG, MB, MN, MT, WF, WJB. Analysis and interpretation of results: TKL, AT, ADJ, SPF, NS, MT, WF, SCS, ER, GP, MBAD, APJ, HRM, CLHH, ANH, SC, WJB. Manuscript preparation: TKL, AT, ADJ, SPF, NS, MT, WF, SCS, ER, VS, CB, GP, MBAD, APJ, HRM, CLHH, ANH, SC, WJB. All authors reviewed the results and approved the final version of the manuscript.

## Data and code availability

The RNA-seq data are deposited in the GEO database, accession number GSE228621. The code used to analyse the data are available from https://github.com/andrewholding/RNASeq-SCN5A.

