## Supplementary Data for "A novel Na_v_1.5-dependent feedback mechanism driving glycolytic acidification in breast cancer metastasis"

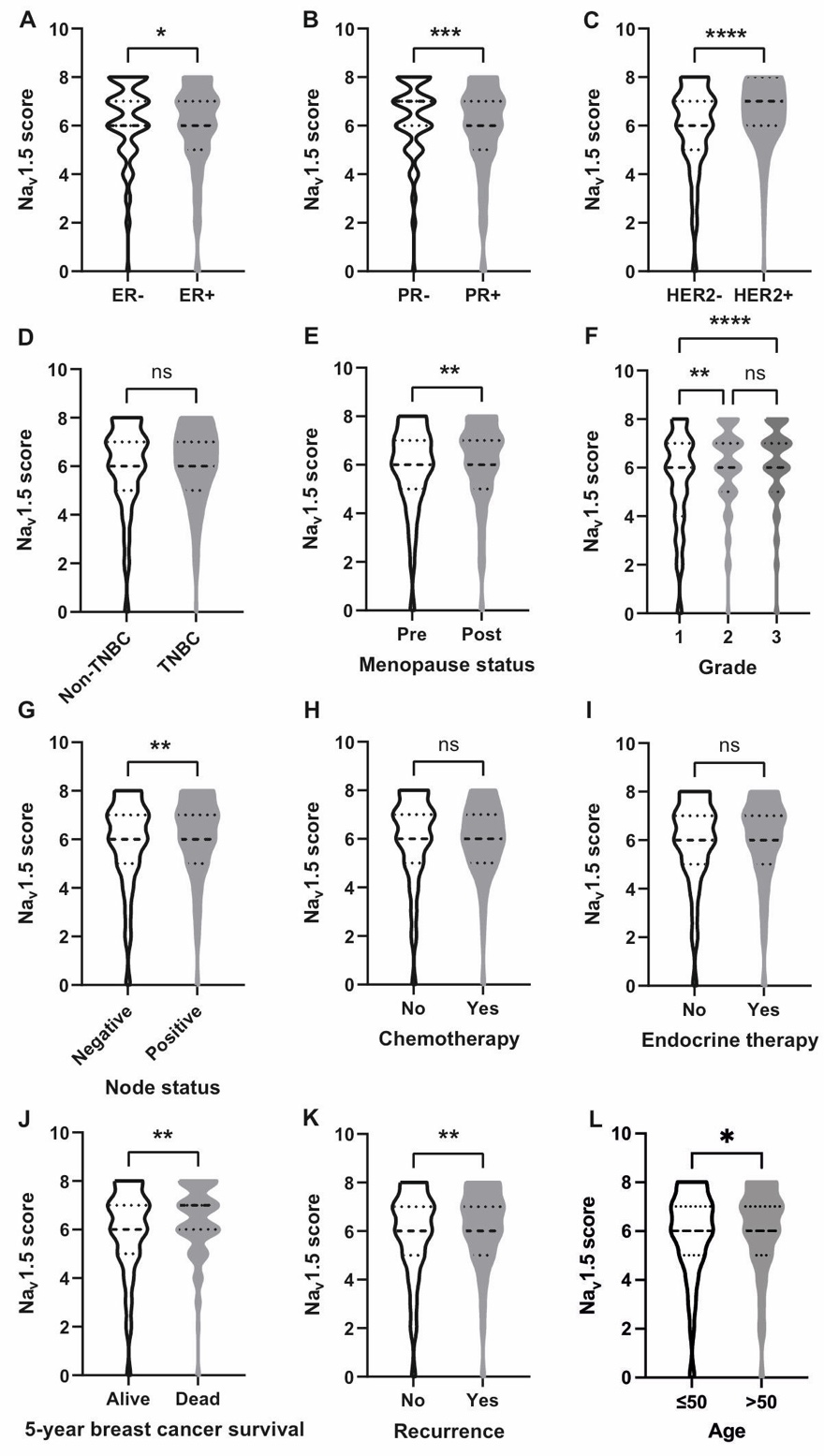

**Supplementary Figure 1.** Na_v_1.5 staining intensity score compared across different histoclinical groups. (A) Estrogen receptor (ER)+ vs. ER-. (B) Progesterone receptor (PR)+ vs. PR-. (C) Human epidermal growth factor 2 (HER2)+ vs. HER2-. (D) Triple-negative breast cancer (TNBC) vs. non-TNBC. (E) Pre- vs. post-menopausal. (F). Tumour grade. (G) Lymph node status. (H) Treated or not treated with chemotherapy. (I) Treated or not treated with endocrine therapy. (J) Five-year survival status. (K) Recurrence status. (L) Age at diagnosis. Data are median ± quartiles. ****P < 0.0001, ***P < 0.001, **P < 0.01, *P < 0.05, ns, not significant. Mann-Whitney tests for all except (F): Kruskal-Wallis with Dunn’s test.

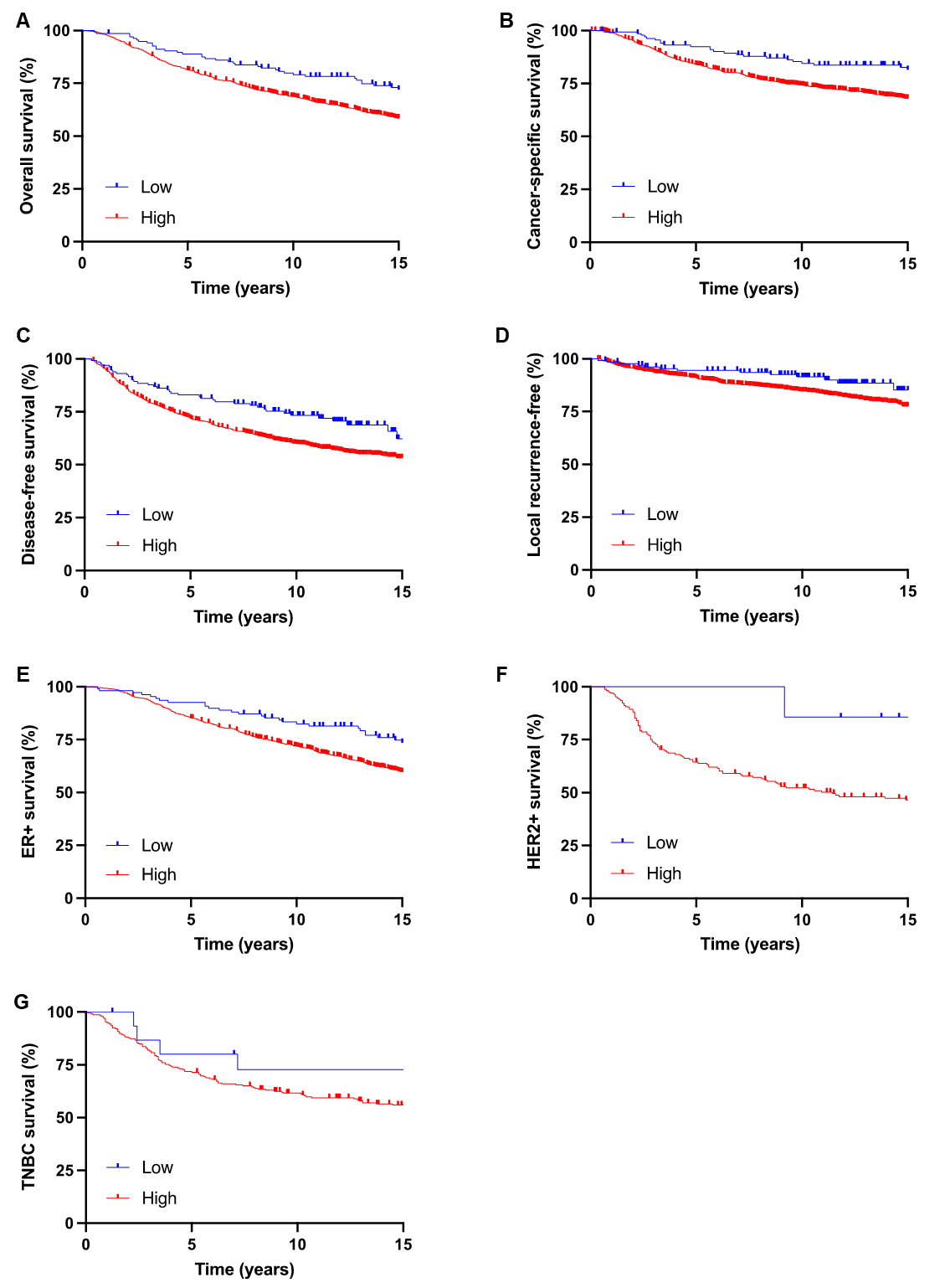

**Supplementary Figure 2.** Survival of breast cancer patients high or low Na_v_1.5-expressing tumours. (A) Overall survival; HR = 1.44 (95% CI 1.12-1.84); P < 0.05 (log-rank test). (B) Cancer-specific survival; HR = 1.87 (95% CI 1.38-2.53); P < 0.01 (log-rank test). (C) Disease-free survival; HR 1.56 (95% CI 1.19-2.04); P < 0.01 (log-rank test). (D) Local recurrence-free survival; HR = 1.86 (95% CI 1.21-2.86); P < 0.05 (log-rank test). (E) Overall survival of patients with estrogen receptor (ER)+ tumours; HR = 1.51 (95% CI 1.14-1.99); P < 0.05 (log-rank test). (F) Overall survival of patients with human epidermal growth factor 2 (HER2)+ tumours; HR = 2.71 (95% CI 1.12-6.52); P = 0.14 (log-rank test). (G) Overall survival of patients with triple-negative breast cancer (TNBC); HR = 1.30 (95% CI 0.62-2.70); P = 0.53 (log-rank test).

**Supplementary Table 1:** Receptor status of patient tumour tissue slices

| Sample | Receptor status |
| --- | --- |
| Patient 1 | ER-  HER2+ |
| Patient 2 | ER+  HER2- |
| Patient 3 | ER+  HER2- |

**Supplementary Table 2:** Characteristics of primary cell samples

| Sample | Type | Receptor status |
| --- | --- | --- |
| 2357N | Normal | n/a |
| 4800N | Normal | n/a |
| 2252N | Normal | n/a |
| 2585N | Normal | n/a |
| 2380T | Tumour | ER+ HER2+ |
| 1691T | Tumour | ER+ HER2+ |
| 2360T | Tumour | TNBC |
| 2285T | Tumour | ER+  HER2+ |
| 1704T | Tumour | HER2+ |
| 1314T | Tumour | ER+ HER2+ |
| 2615T | Tumour | ER+ HER2+ |
| 1958T | Tumour | TNBC |
| 2724T | Tumour | ER+ HER2+ |
| 2599T | Tumour | TNBC |
| 1997T | Tumour | TNBC |
| 2424T | Tumour | TNBC |
| 2753T | Tumour | TNBC |

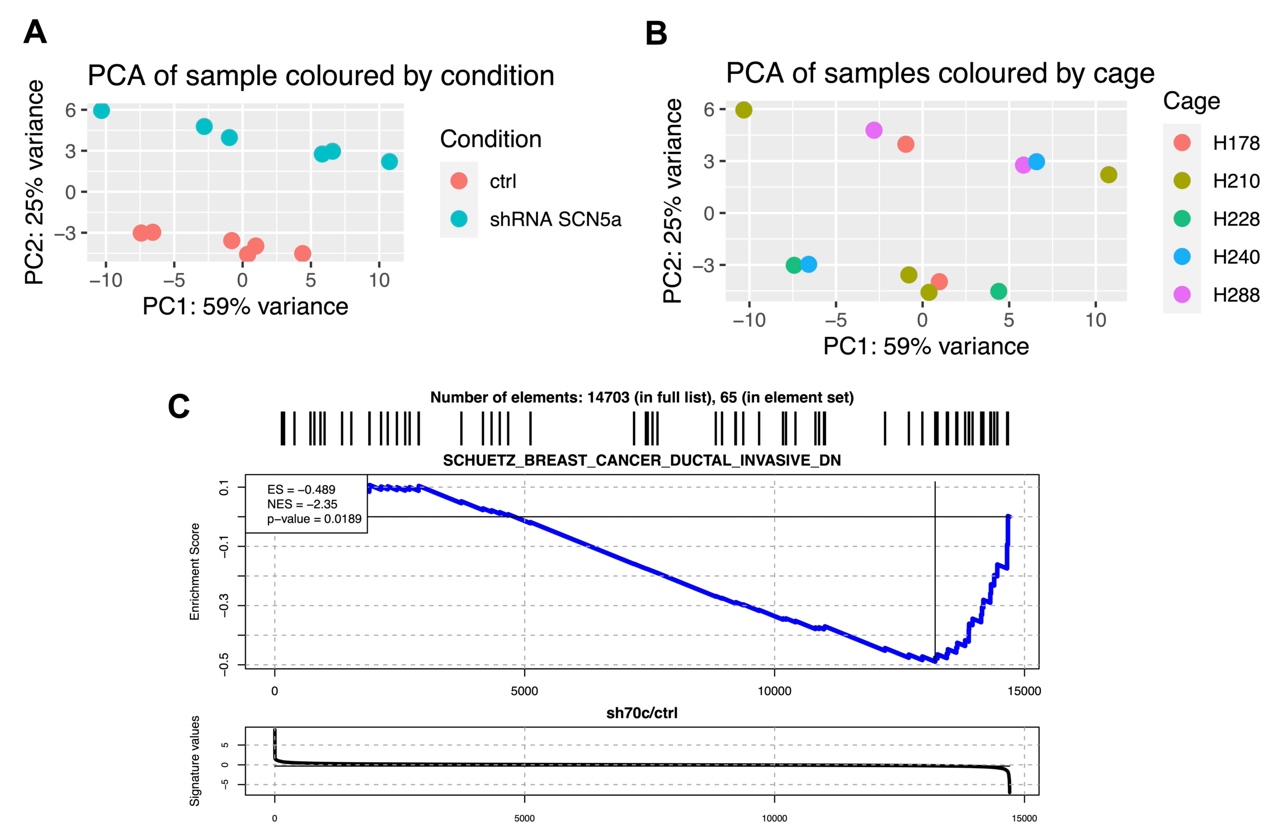

**Supplementary Figure 3.** Transcriptional response to *SCN5A* downregulation in xenograft tumours. (A) Principal component analysis (PCA) biplot of tumour sample coloured by cell type (shControl vs shRNA *SCN5A*). (B) PCA biplot of tumour sample coloured by animal cage. (C) Gene set enrichment analysis (GSEA) of invasion-downregulated genes defined in the MSigDB gene set SCHUETZ_BREAST_CANCER_DUCTAL_INVASIVE_DN analysed in Na_v_1.5 knockdown vs. control tumours.

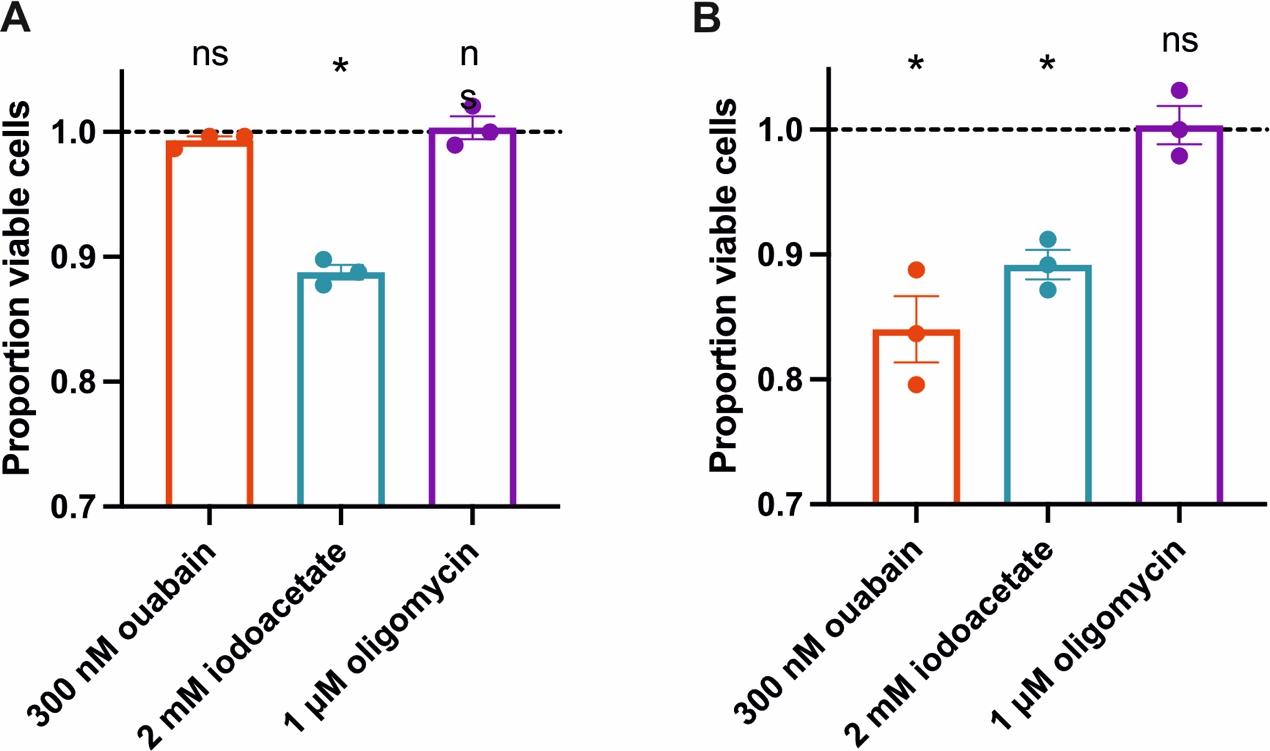

**Supplementary Figure 4.** Effect of glycolytic and mitochondrial respiration on viability of MDA-MB-231 and MCF7 cells. (A) Viability of MDA-MB-231 cells after 6 h treatment with ouabain (300 nM), or 2 h treatment with sodium iodoacetate (2 mM) or oligomycin (1 µM). Data are normalised to the ratio in vehicle-treated cells (n = 3 experimental repeats; one sample *t* tests). (B) Viability of MCF7 cells after 6 h treatment with ouabain (300 nM) or 2 h treatment with sodium iodoacetate (2 mM) or oligomycin (1 µM). Data are normalised to the ratio in vehicle-treated cells (n = 3 experimental repeats; one sample *t* tests). Data are mean ± SEM. *P < 0.05, ns, not significant.

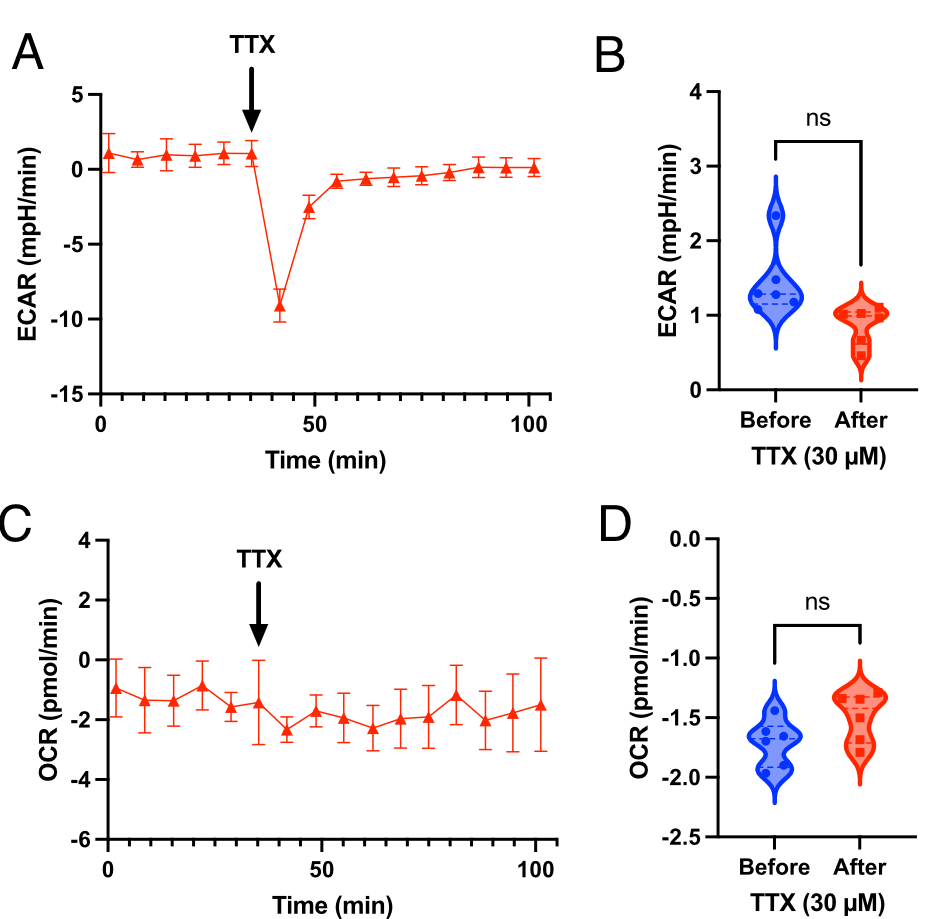

**Supplementary Figure 5.** Effect of TTX citrate buffering on measurements of extracellular acidification rate (ECAR) and oxygen consumption rate (OCR). (A) Representative measurements of ECAR in wells containing medium only (no cells). TTX was added to the treated cells at the indicated timepoint, to give a final concentration of 30 µM (n = 3 wells). (B) ECAR compared before (first 6 timepoints) vs. after TTX addition (last 6 timepoints). (C) Representative measurements of OCR in wells containing medium only. TTX was added to the treated cells at the indicated timepoint, to give a final concentration of 30 µM (n = 3 wells). (D) OCR compared before (first 6 timepoints) and after TTX addition (last 6 timepoints). Data are mean ± SEM. ns, not significant (2-way ANOVA).

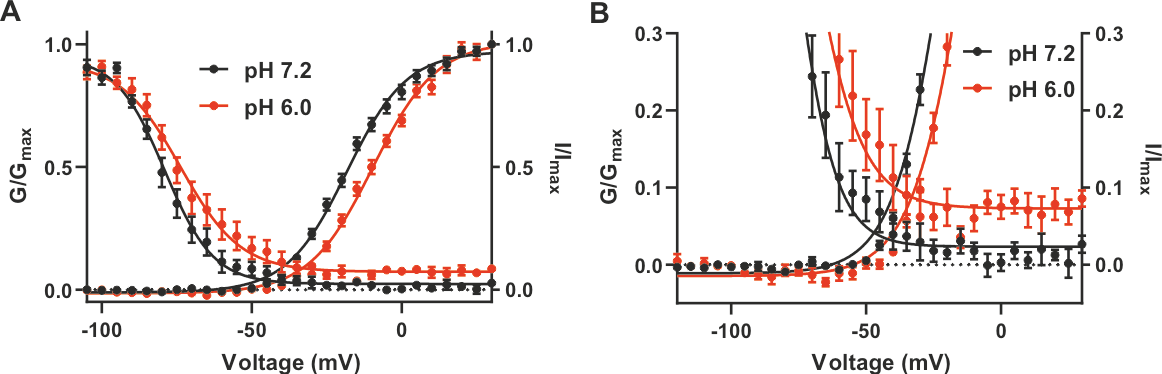

**Supplementary Figure 6.** Effect of pH_e_ 6.0 on VGSC gating in MDA-MB-231 cells. (A) Overlay of activation and inactivation curves at pH 7.2 and 6.0 (n = 8 cells with the largest currents). (B) Expanded data from (A) showing the window current. Data are mean ± SEM.

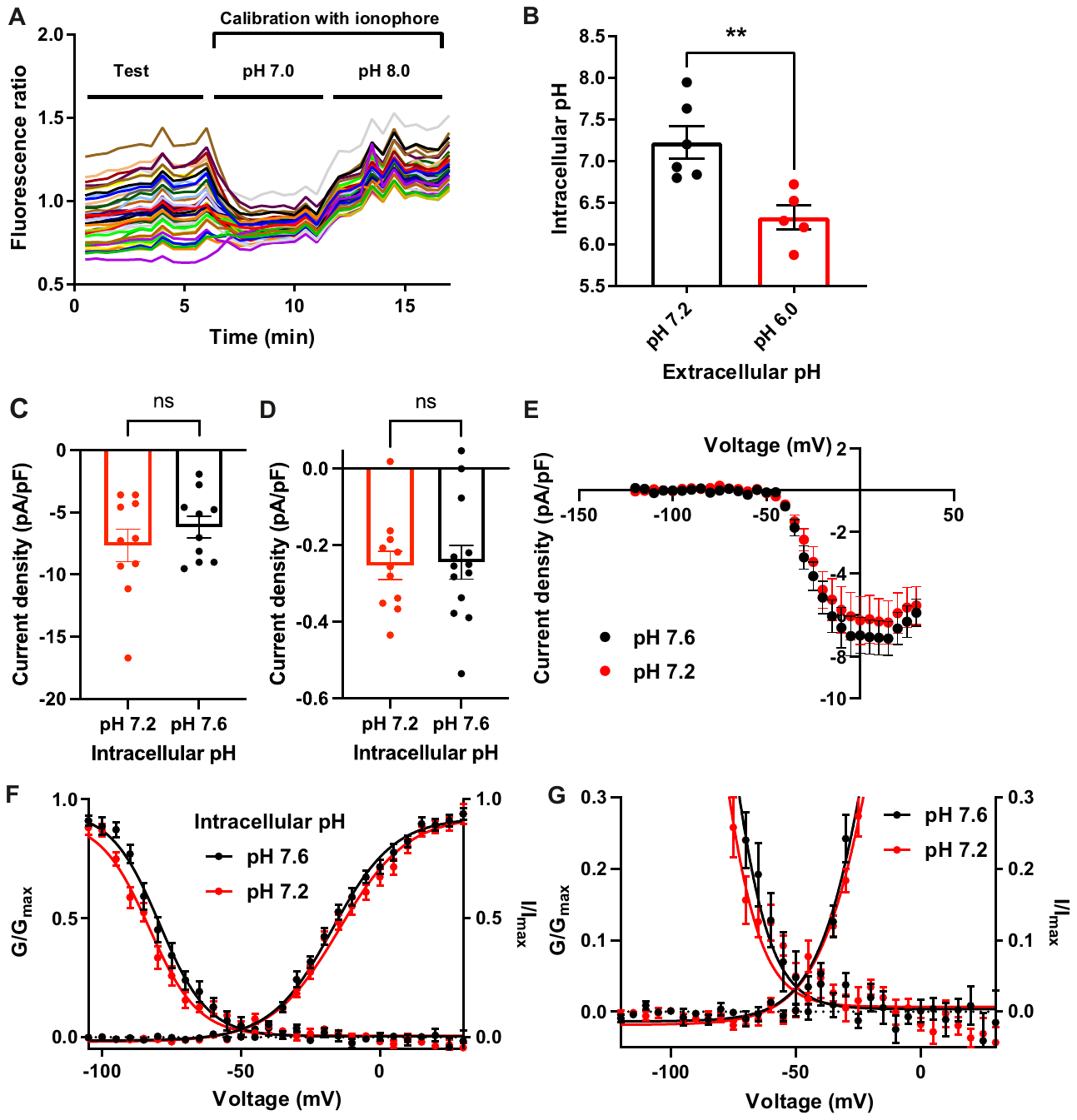

**Supplementary Figure 7.** Effect of intracellular pH on VGSC currents and gating. (A) Example two-point calibration of BCECF-AM fluorescence ratio in MDA-MB-231 cells on a coverslip (each coloured line is one cell). (B) Effect of 10-minute incubation with physiological saline solution at pH 7.2 or 6.0 on pH_i_ in MDA-MB-231 cells (n ≥ 5 experiments of 40 cells each; unpaired *t* test). (C) Peak Na^+^ current density at pH 7.2 and pH 7.6 in patch pipette (n = 10 cells, unpaired *t* test). (D) Mean persistent Na^+^ current density measured 20-25 ms following depolarisation (n = 11-13 cells; unpaired *t* test). (E) Average current density/voltage relationship at pH_i_ 7.2 or 7.6 (n = 10-12 cells). (F) Activation and steady-state inactivation curves at pH_i_ 7.2 and 7.6 (n = 10-12 cells). (G) Expanded graph from (F) showing the window current, which is unchanged between pH_i_ of 7.2 and 7.6. Data are mean ± SEM. **P < 0.01, ns, not significant.

**Calculation to predict Na_v_1.5-dependent extracellular acidification rate**

Assumptions:

Persistent Na^+^ current into breast cancer cells through VGSCs is counteracted by activity of NKA to maintain a stable [Na^+^]_i_. NKA pumps 3 Na^+^ ions out of the cell per cycle in which it hydrolyses one molecule of ATP (Post & Jolly, 1957). Glycolytic production of lactic acid produces 2 molecules of ATP and 2 molecules of lactate per glucose. The production of H^+^ by this reaction coupled to the ATPase hydrolysis of ATP to ADP generates 2 H^+^ per glucose (Hochachka & Mommsen, 1983). There is therefore net production of one H^+^ per ATP molecule generated by glycolytic fermentation to produce lactate.

Therefore:

*(H^+^ evolution rate) = (Na^+^ influx rate)/3	[1]*

It is also assumed that all H^+^ produced is removed via NHE1 to maintain a stable intracellular pH:

*(H^+^ evolution rate) = (NHE1 rate)	[2]*

Since this extra NHE1 activity leads to further Na^+^ entry (1:1 exchange of Na^+^ for H^+^):

*(Na^+^ influx rate) = {(Na_v_1.5 persistent current) / F} + (NHE1 rate)	[3]*

where F = Faraday constant

So at intracellular steady state of [H^+^] and [Na^+^]_i_, substituting in *[3]* from *[1]* and *[2]*:

*3 x (H^+^ evolution rate) = {(Na_v_1.5 persistent current) / F} + (H^+^ evolution rate)*

which is equivalent to:

*(H^+^ evolution rate) = {(Na_v_1.5 persistent current)/(2 x F)}*

Calculating the Na_v_1.5 persistent current

Extracellular acidification through NHE1 due to Na_v_1.5 activity happens in caveolae (Brisson *et al.*, 2013). The shape of these caveolae could provide a localised extracellular microenvironment with a particularly low pH. We showed earlier that in MDA-MB-231 cells at pH_e_ 6.0, the maximal transient Na^+^ current density was -9.19 ± 1.28 pA/pF. The availability of channels at the resting membrane potential of -18.9 mV was 10.3 ± 2.2% of the maximal transient current (Figure 6J) so the persistent current density was -0.95 ± 0.13 pA/pF.

Na_v_1.5 persistent current = -0.95 ± 0.13 pA/pF at pH 6.0

Whole cell capacitance = 26.0 ± 2.2 pF

Cell density for MDA-MB-231 cells in Seahorse analyzer: 3 x 10^4^ / well

F = 96500 C/mol

Therefore:

*Molar (H^+^ evolution rate) per well = (Na_v_1.5 persistent current density) x (Cell Capacitance) x (Cell density) /(2 x F)*

*Molar (H^+^ evolution rate) per well = (0.95 x 10^-12^ x 26 x 3 x 10^4^) / (2 x 96500)*

*= 3.83 x 10^-12^ mol/s*

*Predicted pH change per second = (Molar H^+^ evolution rate) / ( (Well Volume) x (Buffer Capacity) )*

Well volume = 200 x 10^-6^ l

Buffer Capacity = 0.897 x 10^-3^ mol/l/pH (Supplementary Table 3)

*Predicted pH change per second = 3.83 * 10^-12^ / (200 x 10^-6^ x 0.897 x 10^-3^)*

*= 2.13 x 10^-5^ pH/s*

*Predicted pH change per minute = 2.13 * 10^-5^ x 60 = 1.28 x 10^-3^ pH/min*

*= 1.3 mpH/min*

**Supplementary Table 3:** Calculation of buffering capacity of medium used in Seahorse analyzer experiments.

| **Buffer** | **Concentration** | **pKa** | **Ka** | **pH** | **Proton conc** | **Buffer factor** |
| --- | --- | --- | --- | --- | --- | --- |
|  | **M** |  | **M** |  | **M** | **moles/litre/pH** |
| **Citrate** | 2.53E-04 | 3.13 | 7.447E-04 | 7.4 | 3.981E-08 | 3.114E-08 |
|  | 2.53E-04 | 4.76 | 1.734E-05 | 7.4 | 3.981E-08 | 1.332E-06 |
|  | 2.53E-04 | 6.40 | 4.018E-07 | 7.4 | 3.981E-08 | 4.779E-05 |
| **Glutamine** | 2.00E-03 | 2.17 | 6.761E-03 | 7.4 | 3.981E-08 | 2.712E-08 |
|  | 2.00E-03 | 9.13 | 7.413E-10 | 7.4 | 3.981E-08 | 8.266E-05 |
| **Glycine** | 4.00E-04 | 2.34 | 4.571E-03 | 7.4 | 3.981E-08 | 8.023E-09 |
|  | 4.00E-04 | 9.60 | 2.512E-10 | 7.4 | 3.981E-08 | 5.740E-06 |
| **Arginine** | 4.00E-04 | 2.17 | 6.761E-03 | 7.4 | 3.981E-08 | 5.424E-09 |
|  | 4.00E-04 | 9.04 | 9.120E-10 | 7.4 | 3.981E-08 | 2.017E-05 |
|  | 4.00E-04 | 12.48 | 3.311E-13 | 7.4 | 3.981E-08 | 7.662E-09 |
| **Cysteine** | 2.00E-04 | 1.96 | 1.096E-02 | 7.4 | 3.981E-08 | 1.672E-09 |
|  | 2.00E-04 | 10.28 | 5.248E-11 | 7.4 | 3.981E-08 | 6.056E-07 |
|  | 2.00E-04 | 8.18 | 6.607E-09 | 7.4 | 3.981E-08 | 5.623E-05 |
| **Histidine** | 2.00E-04 | 1.82 | 1.514E-02 | 7.4 | 3.981E-08 | 1.211E-09 |
|  | 2.00E-04 | 9.17 | 6.761E-10 | 7.4 | 3.981E-08 | 7.563E-06 |
|  | 2.00E-04 | 6.00 | 1.000E-06 | 7.4 | 3.981E-08 | 1.696E-05 |
| **Isoleucine** | 8.00E-04 | 2.36 | 4.365E-03 | 7.4 | 3.981E-08 | 1.680E-08 |
|  | 8.00E-04 | 9.60 | 2.512E-10 | 7.4 | 3.981E-08 | 1.148E-05 |
| **Leucine** | 8.00E-04 | 2.36 | 4.365E-03 | 7.4 | 3.981E-08 | 1.680E-08 |
|  | 8.00E-04 | 9.60 | 2.512E-10 | 7.4 | 3.981E-08 | 1.148E-05 |
| **Lysine** | 8.00E-04 | 2.18 | 6.607E-03 | 7.4 | 3.981E-08 | 1.110E-08 |
|  | 8.00E-04 | 8.95 | 1.122E-09 | 7.4 | 3.981E-08 | 4.912E-05 |
|  | 8.00E-04 | 10.53 | 2.951E-11 | 7.4 | 3.981E-08 | 1.364E-06 |
| **Methionine** | 2.00E-04 | 2.28 | 5.248E-03 | 7.4 | 3.981E-08 | 3.494E-09 |
|  | 2.00E-04 | 9.21 | 6.166E-10 | 7.4 | 3.981E-08 | 6.918E-06 |
| **Phenylalanine** | 4.00E-04 | 1.83 | 1.479E-02 | 7.4 | 3.981E-08 | 2.479E-09 |
|  | 4.00E-04 | 9.13 | 7.413E-10 | 7.4 | 3.981E-08 | 1.653E-05 |
| **Serine** | 4.00E-04 | 2.21 | 6.166E-03 | 7.4 | 3.981E-08 | 5.948E-09 |
|  | 4.00E-04 | 9.15 | 7.079E-10 | 7.4 | 3.981E-08 | 1.581E-05 |
| **Threonine** | 8.00E-04 | 2.09 | 8.128E-03 | 7.4 | 3.981E-08 | 9.024E-09 |
|  | 8.00E-04 | 9.10 | 7.943E-10 | 7.4 | 3.981E-08 | 3.534E-05 |
| **Tryptophan** | 8.00E-05 | 2.83 | 1.479E-03 | 7.4 | 3.981E-08 | 4.959E-09 |
|  | 8.00E-05 | 9.39 | 4.074E-10 | 7.4 | 3.981E-08 | 1.847E-06 |
| **Valine** | 8.00E-04 | 2.32 | 4.786E-03 | 7.4 | 3.981E-08 | 1.532E-08 |
|  | 8.00E-04 | 9.62 | 2.399E-10 | 7.4 | 3.981E-08 | 1.097E-05 |
| **Pyruvate** | 5.00E-04 | 2.45 | 3.548E-03 | 7.4 | 3.981E-08 | 1.292E-08 |
| **TTX** | 3.00E-05 | 8.76 | 1.738E-09 | 7.4 | 3.981E-08 | 2.769E-06 |
|  |  |  |  | **OrganicSum** | | **4.03E-04** |
| **NaH_2_PO_4_** | 9.00E-04 | 2.16 | 6.918E-03 | 7.4 | 3.981E-08 | 1.193E-08 |
|  | 9.00E-04 | 7.21 | 6.166E-08 | 7.4 | 3.981E-08 | 4.942E-04 |
|  | 9.00E-04 | 12.32 | 4.786E-13 | 7.4 | 3.981E-08 | 2.492E-08 |
|  |  |  |  | **InorganicSum** | | **4.94E-04** |
| **WaterTerm** |  |  |  | 7.4 | 3.981E-08 | 2.910E-07 |
|  |  |  |  | **WaterTerm** | | **2.91E-07** |
|  |  |  |  | **TotalBufferCapacity** | | **8.97E-04** |

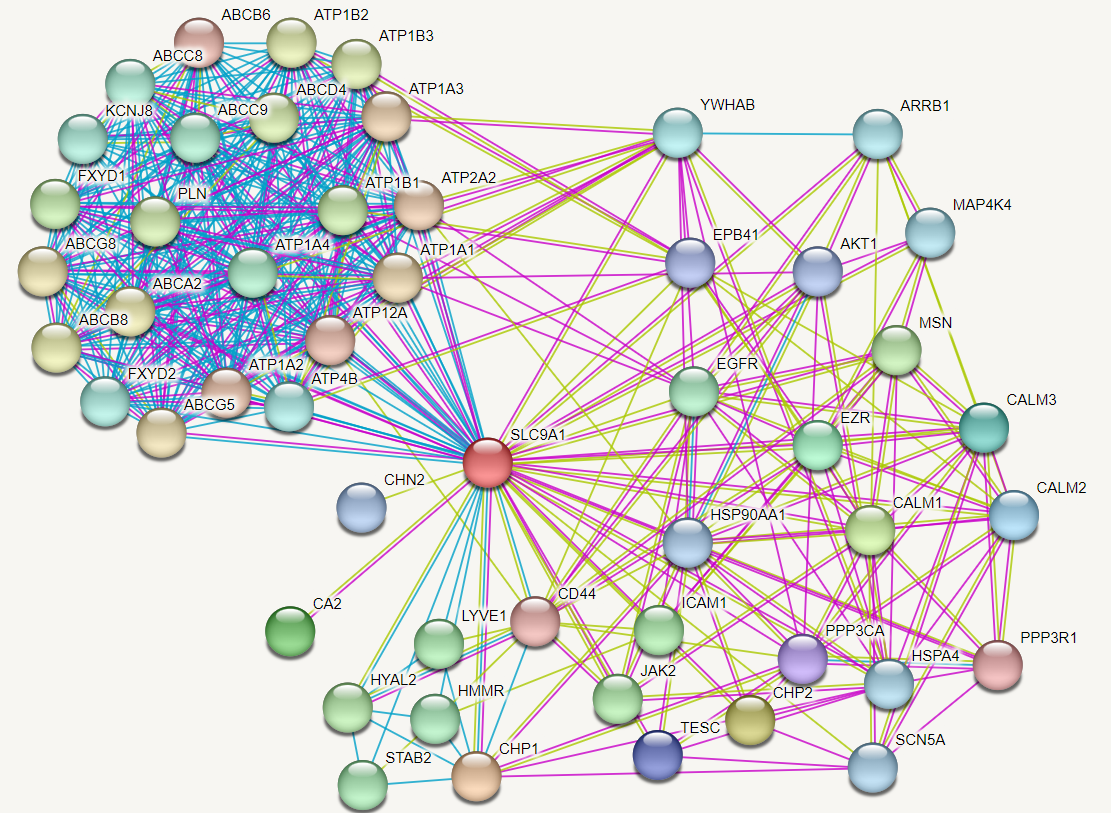

**Supplementary Figure 8**. *In silico* analysis of protein-protein interactions of NHE1 *(SLC9A1)*, created using the STRING database v11.5 © STRING consortium 2022. Only the physical subnetwork is shown (proteins must be part of a physical complex). Evidence for interactions is colour-coded: cyan = from curated databases, magenta = experimentally determined, pale green = text mining. Genes with names *ATP1A** or *ATP1B** are NKA subunits.
